# Reprogramming mRNA localization by targeted RNA-protein interference reveals roles in oncogenesis

**DOI:** 10.64898/2026.02.27.708609

**Authors:** Devon E. Mason, Diptankar Bandyopadhyay, Neal Jiwnani, Brigitte Meyer, Irving Garcia Jimenez, Ramiro Iglesias-Bartolome, Stavroula Mili

## Abstract

RNA binding proteins (RBPs) associate with RNAs in intricate ribonucleoprotein complexes and regulate various aspects of RNA life cycle and, by extension, cell functions. Despite their significance, elucidating the functional contributions of specific RNA-RBP binding events, particularly in long-term phenotypic assays, remains challenging. Here, we harness the specificity of CRISPR/dCas13 to interfere with specific RNA-RBP interactions. We apply this methodology to GA-rich mRNA localization elements which recruit the RNA-binding protein CNBP and serve as platforms for the assembly of mRNA trafficking complexes. We show that dCas13/gRNA binds to target transcripts in a highly specific manner and interferes with CNBP recruitment leading to altered target mRNA localization and cell motility, consistent with the function of the targeted mRNAs. We describe optimizations and considerations for the stable implementation of this system. We further use it to show that interfering with the localization of the GA-containing *NET1* mRNA reduces cancer cell invasiveness and metastatic lung colonization. These data reveal a potential role for mRNA localization in oncogenic progression and illustrate how this CRISPR-based system can be used for long-term functional investigations of RNA-RBP interactions *in vivo*.

**GRAPHICAL ABSTRACT:** 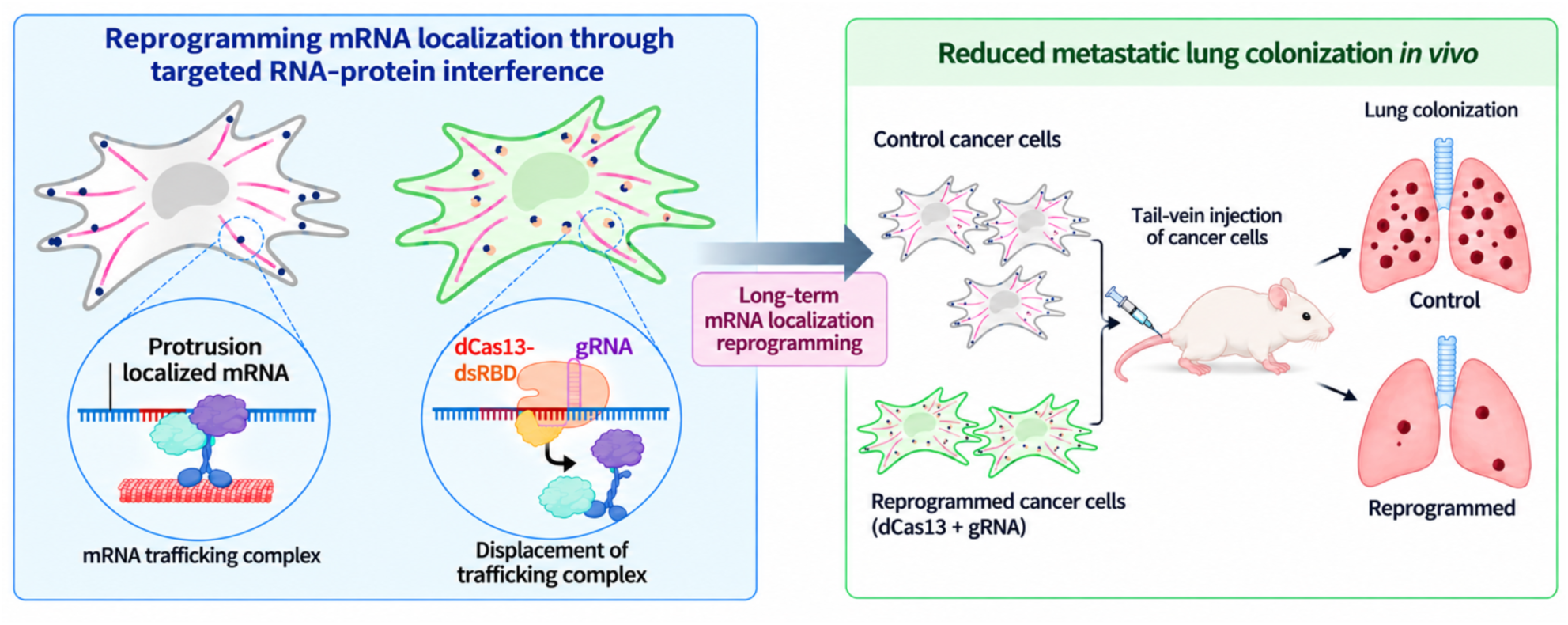

## INTRODUCTION

Post-transcriptional regulation of gene expression is mediated primarily by RNA-binding proteins (RBPs) which dynamically interact with specific sequences or structural motifs within target mRNAs. RNA-RBP binding events affect virtually all aspects of mRNA metabolism including processing, stability, translation and localization [1–3]. The 3’-UTRs of mRNAs often harbor multiple sequence elements that recruit specific RBPs which orchestrate complex spatiotemporal regulation to shape the cellular proteome [4]. Underscoring the importance of these interactions, their mis-regulation is associated with various diseases, including cancer [5–7].

Recent advances in transcriptome-wide mapping of RBP binding sites have substantially enhanced our view of ribonucleoprotein assembly and organization [8]. Nonetheless, studying specific RNA-RBP interactions and understanding their individual functional contributions remains largely challenging. Typical approaches involving depletion of RBPs can have indirect effects since a single RBP can bind multiple RNAs and hence, has a complex regulatory network. On the other hand, precise genetic deletion or mutation of cis-regulatory RNA sequence elements can be cumbersome, is difficult to temporally control, and is irreversible since it involves alteration of the genomic DNA sequence. An alternative approach utilizing antisense oligonucleotides (ASOs) offers a flexible methodology to block specific RNA-RBP interactions [9–13]. However, ASO effects in dividing cells are short-lived and do not allow the study of long-term phenotypic consequences upon sequence specific RNA-RBP perturbation. The above reasons highlight the need to develop a tool that can selectively disrupt sequence-specific RNA-RBP interactions to gain functional insights in disease relevant models and timescales.

Due to the minimal nature and compact organization of the effector protein, Class 2 CRISPR-Cas systems have emerged as important tools for directing nucleic acid manipulation [14,15]. Type VI CRISPR-Cas systems, in particular the RNA-targeting CRISPR-Cas13 based systems, are being widely utilized as a programmable switch for directing a variety of transcript-specific effects including RNA processing, editing, localization as well as mapping transcript specific proteome interactions [16–21]. These approaches exploit the sequence specificity of the guide RNA (gRNA) to recruit catalytically inactive dCas13/gRNA complexes to specific RNA regions wherein the effector enzyme/factor which is fused with the Cas protein exerts its effect on the target RNA.

Here, we sought to exploit the RNA-targeting ability of dCas13 to develop a genetically encoded ASO analog which can sterically compete and disrupt specific RNA-RBP interactions. To benchmark this system, we applied it towards GA-rich 3’UTR elements that mediate mRNA trafficking to peripheral cell protrusions [12,13,22–24]. GA-rich elements direct the formation of an RNA transport complex by recruiting the RNA-binding protein CNBP and the microtubule motor KIF1C [25,26]. ASOs directed against GA-rich elements disrupt formation of the mRNA-CNBP-KIF1C complex and consequently diminish mRNA trafficking, leading to cellular phenotypes of decreased migratory speed and invasiveness [12,13,26–28]. Therefore, GA-rich element-based mRNA regulation offers a biologically relevant setting for assessing and standardizing CRISPR-dCas13/gRNA efficacy as a functional interference tool.

Here, we find that the strength of target RNA binding as well as the amount of cytoplasmic gRNA limit the ability of dCas13/gRNA complex to compete with RNA element function. We describe optimizations that allow the implementation of a fully genetically encoded, stably expressed CRISPR-dCas13/gRNA system that disrupts GA-dependent mRNA localization. This effect is mediated by competition of CNBP binding and is comparable in magnitude with ASO-based interventions leading to robust alterations in cell behaviors. We combine this tool with long-term functional studies, demonstrating that mistargeting of the GA-containing *NET1* mRNA leads to reduced cancer spheroid invasiveness in vitro, and reduced metastatic lung colonization in mice. This work describes a new strategy to harness the specificity of CRISPR-Cas tools for selective and controllable interference of specific RNA-RBP interactions, providing a necessary step towards understanding their long-term functional contributions.

## MATERIALS AND METHODS

### Cell culture

MDA-MB-231 cells (ATCC; HTB-26) were cultured in Leibovitz’s L-15 medium (Gibco; 11415064) supplemented with 10% Fetal Bovine Serum (FBS; Thermo Fisher A31604-02) or tetracycline-free FBS (Thermo Fisher A47362-01) and 1% Penicillin-Streptomycin (Gibco; 15140122) at 37°C with ambient CO_2_ in a humidified incubator. HEK293T cells (ATCC; CRL-3216) were cultured in Dulbecco’s modified eagle medium (DMEM; Gibco; 11995-065) supplemented with 10% FBS and 1% Penicillin-Streptomycin at 37°C and 5% CO_2_ in a humidified incubator. Cells were passaged in 0.05% trypsin (Gibco; 25300054) and were confirmed to be mycoplasma negative by regular PCR testing.

Lentiviral particles used to generate stable cell lines were produced in HEK293T cells. Briefly, lentivectors and packaging plasmids pMD2.VSVG (Addgene plasmid #12259) and psPAX2 (Addgene plasmid #12260) were transfected with PolyJet DNA transfection reagent (SignaGen; SL100688). Lentiviral particle containing media were collected after 48 hours and incubated with Polyethylene Glycol at 4°C overnight prior to concentration by centrifugation. Concentrated particles were used to infect cells, and stably transduced lines were generated after selection for 10 days with construct-specific antibiotics (Puromycin [Gibco; A11138-03], 6 μg/mL Blasticidin [Gibco; A11139-03], or 0.35 mg/mL Geneticin [Gibco; 10131-035]). dCas13-EGFP or Cherry-NLS expressing cells were further selected by fluorescence activated cell sorting (FACS).

### Transfection

For transient transfection experiments, cells were replated 24 hours before transfection of synthetic gRNA using lipofectamine RNAimax (Invitrogen; 13778150) or synthetic gRNA with DNA plasmids using Lipofectamine LTX (Invitrogen; A12621) according to the manufacturer’s instructions. Cells were plated to be ∼70-80% confluent at the time of transfection and were assayed 48-72 hours after transfection. For imaging experiments cells were transfected in a 12 well plate with 0.1 nmol of synthetic gRNA and/or 500ng of plasmid

DNA. For RNA immunoprecipitation experiments transfection reactions were scaled for 6cm dishes and 0.35 nmol of synthetic gRNA was used per dish.

Synthetic gRNAs were designed to contain the processed form of the RfxCas13 direct repeat (30nt; Supplementary Table S1; [29]). Spacer sequences were designed to hybridize to specific target mRNA regions, which were chosen to minimize off targets and not affect mRNA abundance or translation [12,13]. Spacer sequence compatibility with Cas13 binding was performed in silico using the web-based “Cas13 design tool” [30]. Synthetic gRNAs were synthesized by IDT to include phosphorothioate 2’-O-methyl bases on the 5’ and 3’ ends to enhance gRNA stability *in vivo*. See Supplementary Table S1 for gRNA sequences used in this study.

### Plasmid cloning

dCas13 variants were cloned into lentiviral vectors under doxycycline-inducible or constitutive promoters. Briefly, dCas13 fragments were PCR amplified from a previously described construct (Addgene plasmid #154938). dCas13 and EGFP-3xFLAG gblocks were subsequently ligated into pcDNA3.1 vector at BamHI and NotI sites. The indicated dsRBDs (Supplementary Table S2) were then inserted upstream of EGFP. The entirety of these constructs were cloned into the pENTR1A backbone and recombined into the pInducer 20 lentivector (Addgene plasmid #44012) using Gateway LR Clonase II (Thermofisher; 11791020). For cloning the dCas13-B2 variant under a CMV promoter, the dCas1-B2-dsRBD-EGFP-FLAG fragment was restriction digested and ligated into the mammalian lentiviral vector pCDH-CMV-MCS-EF1-Puro (System Biosciences; CD510B-1) into BamHI and NotI sites.

RfxCas13 gRNAs were designed to contain either single processed (30 nt) or unprocessed direct repeats (36 nt) with identical spacers as previously described [29]. See Supplementary Table S1 for gRNA sequences used in this study. To generate a lentiviral expression cassette the pLenti-CMV-TetR (Addgene plasmid #17492) was digested with ClaI and XbaI sites followed by ligation of a human U6 (hU6) promoter fragment upstream of two BsmBI sites and a stretch of thymidines, transcribed as a polyuridine tract for transcription termination. DNA fragments corresponding to individual gRNAs were cloned into BsmBI sites. gRNA arrays were generated using the NEBridge Golden Gate Assembly kit (NEB; E1602S) using PCR-generated gRNA fragments containing unique overlapping overhangs (Supplementary Table S1).

For recombinant CNBP expression and purification, E. coli codon optimized human CNBP sequence was cloned into the bacterial expression vector pET-32a(+) (Novagen; 69015-3). Specifically, gBlock gene fragments containing the sequence encoding the tobacco etch virus (TEV) protease cleavage site followed by human CNBP coding sequence along with BamHI and NotI restriction sequences on either end of the TEV-CNBP sequence were procured from Integrated DNA Technologies. The gBlock fragment was inserted into the pET-32a(+) vector by restriction digestion followed by ligation.

Lentiviral Cherry-NLS constructs were generated as previously described [13].

### Fluorescent in situ hybridization

For fluorescent *in situ* hybridization (FISH) imaging, MDA-MB-231 cells were plated on collagen IV (10 μg/mL; Sigma; C5533) coated coverslips in 0.1% serum containing basal media. Cells were allowed to adhere for 2-3 hours then washed in PBS followed by fixation in 4% paraformaldehyde in PBS for 20 min at room temperature.

For mRNA imaging, FISH was performed using the ViewRNA ISH Cell Assay kit (ThermoFisher; QVC0001) according to the manufacturer’s protocol. The following probes were used: human *NET1* (VA6-3169338), human *RAB13* (VA6-3168453), *GFP* (VF1-14510).

After mRNA labeling, cells were stained with HCS green cell mask (ThermoFisher; H32714) and DAPI to delineate cell and nuclear outlines. Samples were mounted with Prolong Gold antifade (ThermoFisher; P36930) overnight prior to imaging.

For gRNA imaging, the RNAscope plus smRNA-RNA HD assay kit was used (Advanced Cell Diagnostics (ACD); 322780) according to the manufacturer’s directions. Briefly, cells were fixed for 30 minutes then washed in PBS. Endogenous peroxidase activity was depleted by incubating coverslips for 10 minutes in hydrogen peroxide followed by protease digestion using protease III diluted 1:15 in PBS for 10 minutes at room temperature. The following probes were used: NET1_1067_ gRNA (ACD; 1258891-S1), *GFP* mRNA (ACD; 400281-C2). For detection, Opal 570 (Akoya Biosciences; OP-001003) or Opal 690 (OP-001006) dyes were used. Samples were then processed as described above for imaging.

### Imaging

Fixed samples were imaged on either a Nikon Eclipse Ti2-E inverted or a Leica SP8 inverted confocal microscope. The Nikon Ti2-E is equipped with a Yokogawa CSU-X1 spinning disk confocal scanner unit, 40× objective (Plan Fluor 40; Oil; NA = 1.30; WD = 240 μm), NIS-Elements software (5.21.03, Build 1489), and a Hamamatsu ORCA-Fusion BT Gen III back-illuminated sCMOS camera. The Leica SP8 is equipped with a HC PL APO 63x oil CS2 objective.

For assessing migration speed, Cherry-NLS expressing cell lines were tracked by live imaging using an Olympus IX81 inverted microscope equipped with an incubation system (Okolab) which maintained the stage and objective at 37°C for the duration of the experiment. Cells were plated in fluorobrite media (ThermoFisher; A18967-01) for 6 hours prior to imaging using a 10x objective. Cells were imaged every 5 minutes for 8 hours.

### Image analysis

For analysis of mRNA distribution in FISH images, a PDI index was calculated using a previously published custom Matlab script [31]. Briefly, the PDI index is an intensity weighted measure of the distribution of an mRNA population relative to the center of the nucleus, on a per cell basis, using max-intensity projected images. It is also normalized for the size and morphology of each cell on a 2D plane, so that a value of 1 reflects a diffuse distribution, values > 1 are more peripheral and values < 1 more perinuclear.

For analysis of gRNA distribution, sum-intensity projections of FISH images were manually segmented into nuclear and cytoplasmic regions using FIJI. Cytosolic gRNA and whole cell *GFP* intensity (Raw integrated density) were measured on a per cell basis. These data had a lognormal distribution and were transformed using the natural log for statistical analysis.

For random cell migration, Cherry-NLS expressing cell lines were used. Additionally, GFP fluorescence was used to identify cells expressing dCas13. The nuclear Cherry fluorescence signal was used to track cells over time using the FIJI plugin TrackMate. Cells dividing, or which could not be tracked for the duration of the experiment were excluded from final analysis.

### RNA immunoprecipitation

To detect RNAs that co-immunoprecipitate with dCas13 we used magnetic FLAG beads (ThermoFisher; A36798) to precipitate dCas13-dsRBD-EGFP-3xFLAG fusions. Briefly, 50 μL of FLAG beads were pre-blocked in BSA, glycogen, Salmon sperm DNA, and E. Coli tRNA. Cells were washed in PBS then lysed in ice cold 1% NP-40 containing buffer with 50 mM Tris (pH 7.4), 150 mM NaCl, 10 mM MgCl_2_, 10% glycerol, 1x HALT protease inhibitor (ThermoFisher; 1861281), and RNase inhibitor (Promega; N2618). Total lysate was first cleared by centrifugation at 10,000xg then incubated for 1 hour at 4°C with FLAG beads while rotating. Beads were washed 5 times in lysis buffer then protein was eluted with 50 μg of FLAG peptide (ThermoFisher; A36806) for 15 minutes at RT with shaking. Total and eluted protein and RNA were further analyzed by western blot, ddPCR, or nanoString nCounter.

### RNA isolation and analysis

For gRNA analysis, RNA was isolated using Trizol LS (Thermo Fisher Scientific; 10296010). The aqueous phase was isolated according to the manufacturer’s instructions. and applied to the Zymo total RNA clean and concentrator kit (Zymo research; R1014) with on-column DNase digestion. For experiments requiring only mRNA isolation the RNeasy Plus Mini Kit (Qiagen; 74134) was used.

RNA was reverse transcribed using the iScript cDNA Synthesis Kit (Bio-Rad; 1768891) and used for droplet digital polymerase chain reaction (ddPCR). PCR reactions were prepared with cDNA, gene specific primers, and ddPCR EvaGreen Supermix (Bio-Rad; 186-4034). Droplets were generated from this reaction using the Automated Droplet Generator (Bio-Rad; 186-4101), PCR was then performed on a C1000 Touch Thermal Cycler (Bio-Rad; 185-1197), and droplet reading was performed on QX-200 Droplet Reader (Bio-Rad; 186-4003). Results were quantified using the QuantaSoft software (Bio-Rad).

For nanoString analysis, immunoprecipated RNA was analyzed using a custom codeset (Supplementary Table S3) and the nCounter analysis system according to the manufacturer’s instructions.

For RNA-seq analysis, total RNA samples were submitted to Plasmidsaurus, Inc. for RNA sequencing. Three biological replicates were analyzed for each sample. Libraries were prepared after polyA selection using a 3′-end counting approach, and Illumina sequenced to generate ∼90-bp single-end reads. Sequencing data were processed by Plasmidsaurus using their standard pipeline, including read quality filtering, alignment to the reference genome, UMI-based deduplication, and gene-level quantification.

### Northern blot

Heat denatured, total cellular RNA was electrophoresed on a 10% Urea-Polyacrylamide gel followed by semi-dry transfer onto a positively charged nylon membrane (Cytiva; RPN119B) for 45 mins at 400mA. Membrane was then dried, UV-crosslinked and prehybridized with Oligo Hybridization Buffer (ThermoFisher; AM8663) at 30^0^C for 60 mins. 5’ IRDye labeled probes (IDT) against U6 snRNA and NET1 gRNA were hybridized at 30^0^C, overnight. A pool of two different NET1 gRNA probes were used that were designed against either the spacer region or the direct repeat region. After sequential washing in 2xSSC/0.1%SDS buffer and 1xSSC/0.1%SDS buffer, blots were developed using an Odyssey fluorescent scanner (Li-Cor).

### Protein production and purification

*Escherichia coli* BL-21 (DE3) cells were transformed with pET-32-TEV-HsCNBP plasmid and single colonies were cultured at 37^0^C in Luria-Bertani (LB) media supplemented with 100ug/ml ampicillin and 0.3mM zinc chloride. Cultures at OD_600nm_ of ∼0.5 were induced with 0.5mM IPTG overnight at 18^0^C. Cells were harvested by centrifugation and pellets were resuspended in lysis buffer (50 mM Tris–HCl pH 7.5; 300 mM NaCl; 10 mM Imidazole; 10 mM β-mercaptoethanol; 10% Glycerol and 1% Protease Inhibitor Cocktail). Cells were lysed using a microfluidizer and the lysate was clarified by centrifugation at 19,000rpm for 30mins at 4^0^C yielding soluble proteins in the supernatant.

Recombinant human CNBP was purified as described in [32] with minor modifications. Briefly, the supernatant expressing Trx-His_6_-TEV-CNBP was initially passed through a 0.5 um filter and the filtrate containing the recombinant protein was allowed to bind slowly to a Ni-NTA chromatography column that was pre-equilibrated with Buffer A (50 mM Tris–HCl pH 7.5; 300 mM NaCl; 30 mM Imidazole; 10% Glycerol and 10 mM β-mercaptoethanol). The column was washed with 5 column volumes of 6 % Buffer B (50 mM Tris–HCl pH 7.5; 300 mM NaCl; 500 mM Imidazole; 10% Glycerol and 10 mM β-mercaptoethanol) in Buffer A and then with 8% Buffer B in Buffer A. Eventually the protein was eluted with 100% Buffer B. The fractions enriched for the protein were identified from the chromatogram peak and confirmed by Coomassie staining of SDS-PAGE gels. These fractions were pooled together and cleaved with TEV protease and dialyzed overnight in Dialysis Buffer A (50 mM Tris–HCl pH 7.5; 300 mM NaCl; 1 mM DTT and 0.1 mM ZnCl_2_). The cleaved, dialyzed product was allowed to batch bind to Ni-NTA agarose resin for 3h at 4^0^C with rotation. The flow-through containing the cleaved, untagged CNBP was collected.

To obtain nucleic acid free-CNBP, the above collected CNBP was further dialyzed in Dialysis Buffer II (50 mM Tris–HCl pH 6.5; 10% Glycerol and 5 μM ZnCl_2_), filtered through a 0.5um membrane and allowed to bind slowly to a Q-Sepharose chromatography column that had been equilibrated with Buffer C (50 mM Tris–HCl pH 6.5; 10% Glycerol and 5 μM ZnCl_2_). Stepwise elution of the protein was done with 100mM, 200mM and 500mM of NaCl in Buffer C. All steps of the purification were carried out at 4^0^C.

### In vitro transcription and RNA isolation

In vitro transcription was performed as described in [26]. Briefly, linearized plasmid templates having the RAB13-3’UTR-BoxB sequence were in-vitro transcribed (4ug per 50ul reaction mix) using T7 Polymerase HC (Promega; P2075) in transcription optimized buffer (Promega) supplemented with 20mM DTT, 10mM MgCl2, 4mM NTPs, 40U/ul RNasin Plus (Promega; N2615) and 1% Triton X-100. Reaction was incubated for 3.5hrs at 37^0^C. Post reaction, the in vitro transcribed RNA was isolated using the Zymo total RNA clean and concentrator kit (R1014). In column DNase digestion was performed to remove plasmid template from the in vitro transcribed RNA.

### In vitro RNA-RBP binding

For purification of dCas13-B2-EGFP-3xFLAG/sgRNA complex, initially, dCas13-B2-EGFP-3xFLAG was immunoprecipitated with anti-FLAG magnetic agarose beads (ThermoFisher; A36798), as described earlier, from cell lysates of HEK 293LT that were transiently expressing the protein. The bound protein was then incubated with molar excess of either non-targeting control (NTC2) or RAB13_191/230_ synthetic gRNA for 15mins at room temperature with shaking. Subsequently, beads were washed 5 times with lysis buffer to remove unbound gRNA, and elution was done with 50ug FLAG peptide (ThermoFisher; A36806) for 15mins with shaking at room temperature to obtain dCas13-B2-EGFP-3xFLAG/sgRNA complex.

For RNA-RBP binding, 850ng of the in-vitro transcribed RNA was incubated with 10ugr of purified λN-GST protein for 15mins at room temperature. The complex was subsequently incubated with 15uL Glutathione magnetic beads (Invitrogen; 78602) for 20 min at room temperature with shaking. The bound RNA-λN-GST protein complex was washed with lysis buffer (50mM Tris-Cl pH 7.5, 0.5% Triton X-100, 100mM NaCl, 2.5mM MgCl_2_) and blocked with lysis buffer containing E. coli tRNA, BSA (Sigma; 10711454001), salmon sperm DNA (Invitrogen; 15632011) and glycogen (Invitrogen; AM9510) for 30–60 min at 4°C. The complex was then incubated with recombinant CNBP and dCas13-B2-EGFP-3xFLAG/sgRNA complex for 1hr at 4^0^C with shaking. Beads were then washed with lysis buffer followed by PBS (pH 8.0) wash and elution with 50ul of 40mM reduced L-Glutathione in PBS (pH 8.0). The eluate was analyzed by Western blot to assess the amount of CNBP and dCas13-B2-EGFP-3xFLAG that associated with the RAB13-3’UTR-BoxB RNA.

### Western blot

For western blotting the following primary antibodies were used: rabbit anti-FLAG (1:2,000; Cell Signaling Technology; A303-138A), rabbit anti-GFP (1:3,000; Abcam; ab11122), mouse anti-α-tubulin (1:3,000; Sigma; T6199), mouse anti-CNBP (1:3,000; Proteintech; 67109), Phospho-eIF2α (1:1000; Cell Signaling Technology; #9721S), eIF2α (1:1000; Cell Signaling Technology; #9722S). Donkey anti-rabbit (800CW; 926-32213) and Donkey anti-mouse (680RD; 926-68072) secondary antibodies from Li-Cor were used at 1:10,000. Membranes were scanned using an Odyssey fluorescent scanner (Li-Cor) and bands were quantified using ImageStudioLite (Li-Cor).

### Cell growth analysis

Cell proliferation was monitored using an Incucyte live-cell analysis system (Sartorius). Cells were seeded at equal densities and imaged at regular intervals under standard cell culture conditions. Cell growth was quantified from phase-contrast images by measuring cell confluence over time using the Incucyte analysis software.

### Spheroid formation and outgrowth assay

For spheroid formation, a single-cell suspension of 0.2 x 10^4^ cells was mixed with Matrigel (Corning, cat# 354234) and placed as a 30ul droplet on a chambered coverglass precoated with Matrigel. After a 30 min incubation at 37^0^C, L15 media supplemented with 10%FBS and 1% Pen-Strep were added. Individual embedded cells were allowed to grow for 8-9 days to form spheroids. To induce invasion, Matrigel-embedded spheroids were gently washed with PBS followed by addition of low serum media (0.1% FBS) and overnight incubation. For imaging, spheroids were stained with 5µM calcein dye (Thermo Fischer Scientific, cat# C3100MP) and 5µM DRAQ5 (Novus Biologicals, cat# NBP2-81125) in PBS for 20 mins at 37^0^C. Spheroids were then fixed with 4%PFA in PBS for 15 mins and imaged on a Leica SP8 inverted confocal microscope using a 10x dry objective. Z-stack images were acquired through the spheroid volume. The total nuclear volume per spheroid, based on DRAQ5 staining, was measured using Imaris (Oxford Instruments). To assess spheroid invasiveness, maximum intensity projections of the z-stacks were used, and a complexity value was calculated, defined as Complexity = Perimeter^2^/(4π x Area).

### Lung colonization model

Mouse studies followed approved protocols by the NIH Intramural Animal Care and Use Committee of the National Cancer Institute. NOD scid gamma mice were purchased from Jackson Laboratory (Strain #:005557). Only female mice were used to match the sex of the MDA-MB-231 breast cancer cells used. 12.7 week old mice were injected intravenously in the tail vein with the corresponding cell line, 5×10e5 cells in 100 μL PBS per mouse. 5 weeks after injection lungs were harvested, perfused and fixed with Z-Fix. For histological analysis, tissues were embedded in paraffin then sectioned (3-μm) for staining by immunofluorescence or HCE. Briefly, paraffin sections were baked for 30 minutes at 60^0^C then deparaffinized using SafeClear II (Fisher Scientific) and ethanol. Rehydrated sections were used for antigen retrieval using the RNAscope Target Retrieval buffer (ACD; 322001) at 60^0^C overnight. After 3x2min washes in PBST, sections were incubated overnight at 4^0^C with Rabbit anti-GFP (1:400; Invitrogen; A-11122) diluted in 3% BSA in PBS. After washing 3x2minutes washes in PBST samples were incubated with Donkey anti-Rabbit Alexa fluor 647 (1:500; Invitrogen; A-31573), Wheat Germ Agglutin Alexa fluor 488 (Invitrogen; W11261; 10 μg/mL), and DAPI for 2 hours at RT. After washing samples were mounted in Prolong gold prior to imaging. The correspondence of GFP-positive areas to tumor tissue was confirmed by HCE staining of adjacent sections.

Histological sections were imaged using either a NanoZoomer Digital Slide Scanner (Hamamatsu) or an Axioscan.Z1 (Carl Zeiss). Adjacent sections were scanned in full using a Plan-Apochromat 20x (NA = 0.8; Carl Zeiss) and imported into the digital pathology platform HALO (Indica Labs). The MiniNet classifier was trained on manually annotated regions to discern GFP+ lesions from normal tissue in multiple slides. Tumor burden was calculated as the ratio of GFP+ over normal tissue area on a per animal basis. Tumor burden and GFP intensity were averaged between adjacent tissue sections for each animal, to compensate for variability in sectioning or staining.

### Statistics and reproducibility

Data are shown as mean ± standard error of the mean (SEM). In Figure 4E and 5B natural log transformed data are displayed. For statistical comparisons parametric tests were used to evaluate significant differences in normally distributed, homoscedastic datasets. Where either of these assumptions were not true, we instead performed non-parametric tests suitable for the data distribution. Specific statistical tests for each experiment are described in the figure captions and p-values were calculated with Prism software package (GraphPad; Version 10.6.1).

## RESULTS AND DISCUSSION

### dCas13-dsRBD fusions augment dCas13-mRNA binding and interfere with GA-rich RNA element function

To assess the ability of CRISPR-dCas13 to interfere with RBP-RNA interactions, we stably expressed a catalytically inactive RfxCas13d protein (dRfxCas13d; hereafter referred to as dCas13) under a doxycycline-inducible promoter. dCas13 was fused to C-terminal eGFP and FLAG tags for biochemical purification and characterization (Fig. 1A). Additionally, a double-stranded RNA-binding domain (dsRBD) was included, to reinforce dCas13/gRNA binding to a target mRNA [21]. We used dsRBDs from various proteins (PKR, PRKRA, DIP1, B2), with differing affinities for double-stranded RNA (Fig. 1A) [33–36]. Doxycycline-inducible dCas13-dsRBD expression cassettes were stably integrated into MDA-MB-231 cells. Transgene induction led to similar expression levels between lines, with the exception of dCas13-DIP1 which consistently exhibited a lower overall expression (Supplementary Fig. S1A). To target dCas13 to specific mRNAs we transfected synthetic guide RNAs (gRNAs) into doxycycline induced cells. gRNAs were designed to target 3’UTR sequences involved in mRNA regulation, or against sequences absent from the human transcriptome (Non-Targeting Control (NTC1, NTC2) gRNAs).

**Figure 1.**
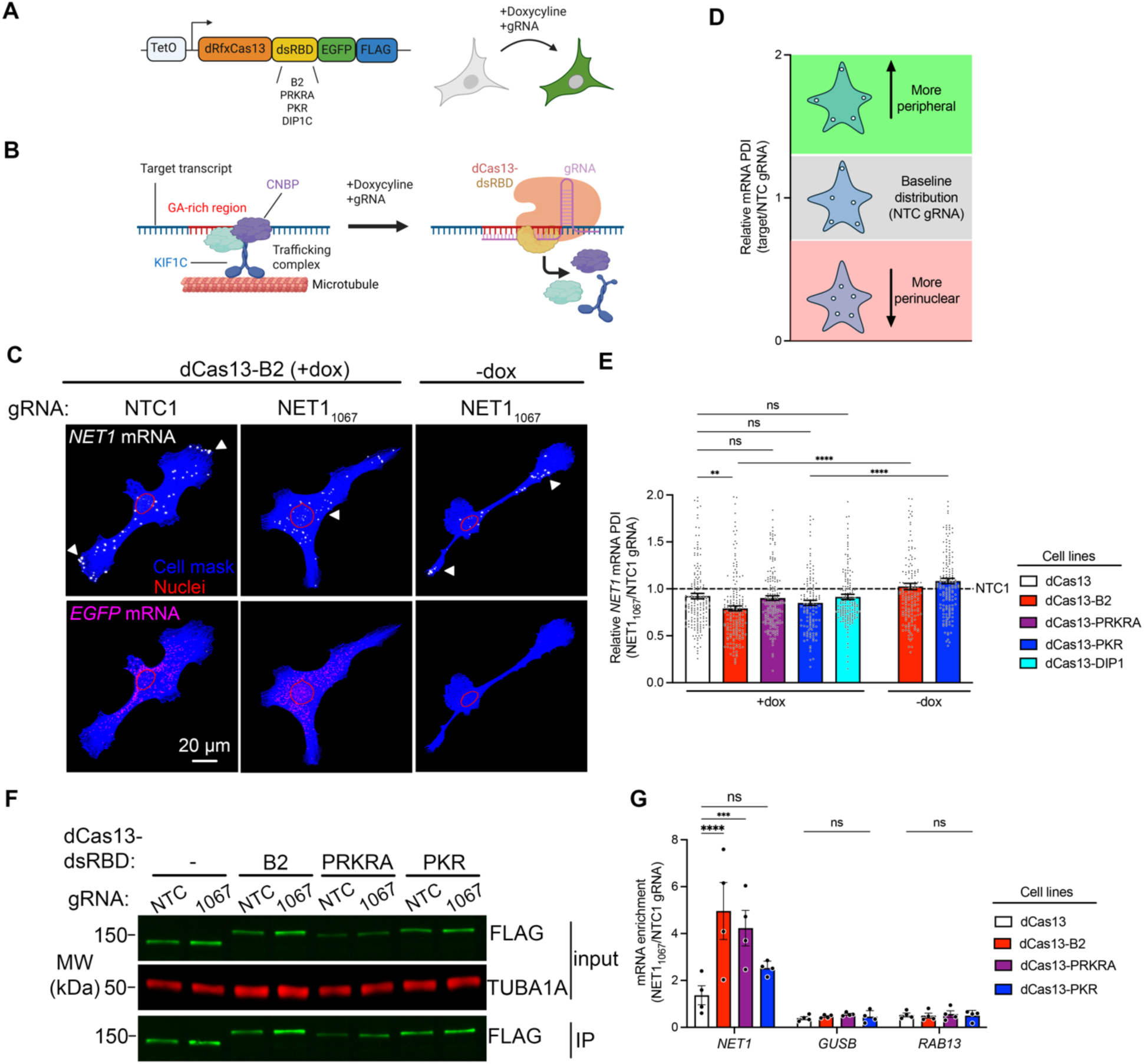
dCas13-dsRBD fusions alter mRNA localization to cell protrusions through enhanced on-target mRNA binding. (**A**) Schematic illustrating the inducible dCas13-dsRBD constructs screened. dCas13-dsRBD expression in MDA-MB-231 cells was induced by doxycycline together with transfection of on-target or non-targeting control (NTC) synthetic gRNAs. (**B**) Schematic depicting the RNA-trafficking complex - consisting of CNBP, KIF1C and putative additional factors - assembled on GA-rich RNA elements. Binding of dCas13-dsRBD/gRNA complex to GA-rich regions is hypothesized to interfere with RBP binding and to lead to trafficking complex disassembly. Created in BioRender. Mason, D. (2026) https://BioRender.com/kdmxt2k (**C**) Representative FISH images of *NET1* and *GFP* mRNAs in dCas13-B2 expressing (+dox) and non-expressing (-dox) cells transfected with the indicated gRNAs. Images of additional conditions are in Supplementary Fig. S1B. Arrowheads point to areas of accumulation of *NET1* mRNA. (**D**) Schematic depicting range of normalized PDI values (target/NTC gRNA) and the corresponding mRNA distributions for GA-containing mRNAs. (**E**) Normalized *NET1* mRNA PDI values in dCas13-dsRBD expressing cells. n = 128-181 cells per condition from 4-6 independent experiments. ** *P* < 0.002, **** *P* < 0.0001, ns: non-significant by Kruskal-Wallis ANOVA with Dunnett’s multiple comparison test. Raw PDI values are in Supplementary Figure S1D. (**F**) Representative western blot images of dCas13-dsRBD immunoprecipitation (IP) using anti-FLAG beads. (**G**) Enrichment of on-target (*NET1*) and off-target (*GUSB* and *RAB13*) mRNAs in dCas13-dsRBD IPs. RNAs were detected by ddPCR and amounts in the presence of NET1_1067_ gRNA were normalized to corresponding NTC gRNA. n = 4 independent experiments per condition. *** *P* < 0.001 and **** *P* < 0.0001 by two-way ANOVA with Dunnett’s multiple comparison test. Bar graphs in panel (E) and (G) represent mean ± SEM with an overlay of individual data points.

To determine whether dCas13-dsRBD fusions can functionally interfere with RBP-RNA interactions, we focused on GA-rich elements, which promote RNA trafficking to the cell periphery through recruiting the RNA-binding protein CNBP and the microtubule motor KIF1C (Fig. 1B) [25,26]. Antisense oligos (ASOs) against GA-rich regions within the *NET1* or *RAB13* mRNAs prevent CNBP binding, leading to more perinuclear distribution of the targeted mRNAs and functional disruption of cell migration [12,13,26,28]. To assess whether CRISPR-dCas13 can similarly interfere with CNBP-dependent trafficking, we designed gRNA to the GA-rich region of the *NET1* mRNA (NET1_1067_ gRNA). Endogenous *NET1* mRNA distribution was assayed by visualizing transcripts using fluorescence in situ hybridization (FISH) (Fig. 1C and Supplementary Fig. S1B). To control for variation in dCas13-dsRBD expression we selected cells with comparable *GFP* RNA amount (Fig. 1C and Supplementary Fig. S1B and C). *NET1* mRNA distribution was quantified by measuring a previously described peripheral distribution index (PDI) [31]. PDI values are normalized against a hypothetical fully diffuse RNA (PDI=1), so that higher PDI values indicate a more peripheral RNA distribution, while lower values denote a perinuclear bias. Supplementary Figure S1D shows the raw *NET1* mRNA PDI values. Some variability in the basal *NET1* mRNA PDI is observed even under control conditions (Supplementary Fig. S1D, see NTC1 gRNA samples), likely due to cell line-specific variation upon stable expression of the various dCas13-dsRBD fusions. To highlight the specific effect of the NET1_1067_ gRNA in each dCas13-dsRBD line, *NET1* PDI values were normalized to those upon NTC1 gRNA delivery (Fig. 1D and E). Interestingly, in the absence of a dsRBD, dCas13/NET1_1067_ gRNA did not appreciably alter *NET1* distribution (Fig. 1E and Supplementary Figure S1D, dCas13 lanes). Instead, dCas13-dsRBD fusions, with the exception of dCas13-DIP1, significantly perturbed *NET1* localization (Supplementary Fig. S1D). The B2 dsRBD had the most pronounced effect on localization and was the only dsRBD to significantly impair localization relative to unmodified dCas13 (Fig. 1E). To exclude the possibility that this effect is attributable to gRNA transfection alone we measured *NET1* PDI in dCas13-B2 and dCas13-PKR cell lines transfected with NET1_1067_ gRNA but without doxycycline induction. In these conditions, *NET1* PDI was unaffected, demonstrating that this effect depends on expression of dCas13 (Fig. 1E and Supplementary Fig. 1D, -dox lanes). Therefore, fusion of dCas13 with a dsRBD augments its ability to interfere with the function of a GA-rich RNA element.

We hypothesized that the increased ability of dCas13-dsRBD fusions to interfere with *NET1* localization stems from their more stable binding with the target mRNA. To test this, we co-immunoprecipitated dCas13 with stably-bound RNAs. Anti-FLAG tag antibodies were used to immunoprecipitate dCas13 fusions in the presence of NTC or NET1_1067_ gRNAs followed by detection of endogenous mRNAs by ddPCR (Fig. 1F and G). Indeed, dsRBD fusions exhibited a higher enrichment of the target *NET1* mRNA compared to dCas13 alone. The B2 dsRBD showed the highest enrichment, in line with its stronger effect on the localization of the *NET1* mRNA shown above. Furthermore, other non-targeted mRNAs (*GUSB, RAB13*) were not appreciably enriched (Fig. 1G) indicating that dCas13 fusions associate specifically with the gRNA-targeted transcript. We conclude that, among the tested dsRBDs, the B2 dsRBD offers an optimal way to stabilize dCas13/gRNA association with a target mRNA and functionally interfere with the targeted sequence. We thus selected dCas13-B2 for further characterization.

### dCas13-B2 interferes with RNA element function in a sequence-specific manner with minimal off-target binding

To assess whether dCas13-B2 can broadly associate with and interfere with functional RNA elements, we designed gRNAs targeting various regions along the *NET1* and *RAB13* 3’UTRs (Fig. 2 A and B). NET1_921_ and NET1_975_ gRNAs target a GA-rich region within the *NET1* 3’UTR, which has been shown to be functionally important for peripheral *NET1* localization, like the region targeted by the NET1_1067_ gRNA. On the other hand, NET1_1351_ gRNA targets a region that is not implicated in *NET1* trafficking [12]. Similarly, RAB13_191_ and RAB13_230_ gRNAs target a GA-rich region important for *RAB13* localization, while RAB13_360_ and RAB13_391_ gRNAs target a non-functional UTR region [13].

**Figure 2.**
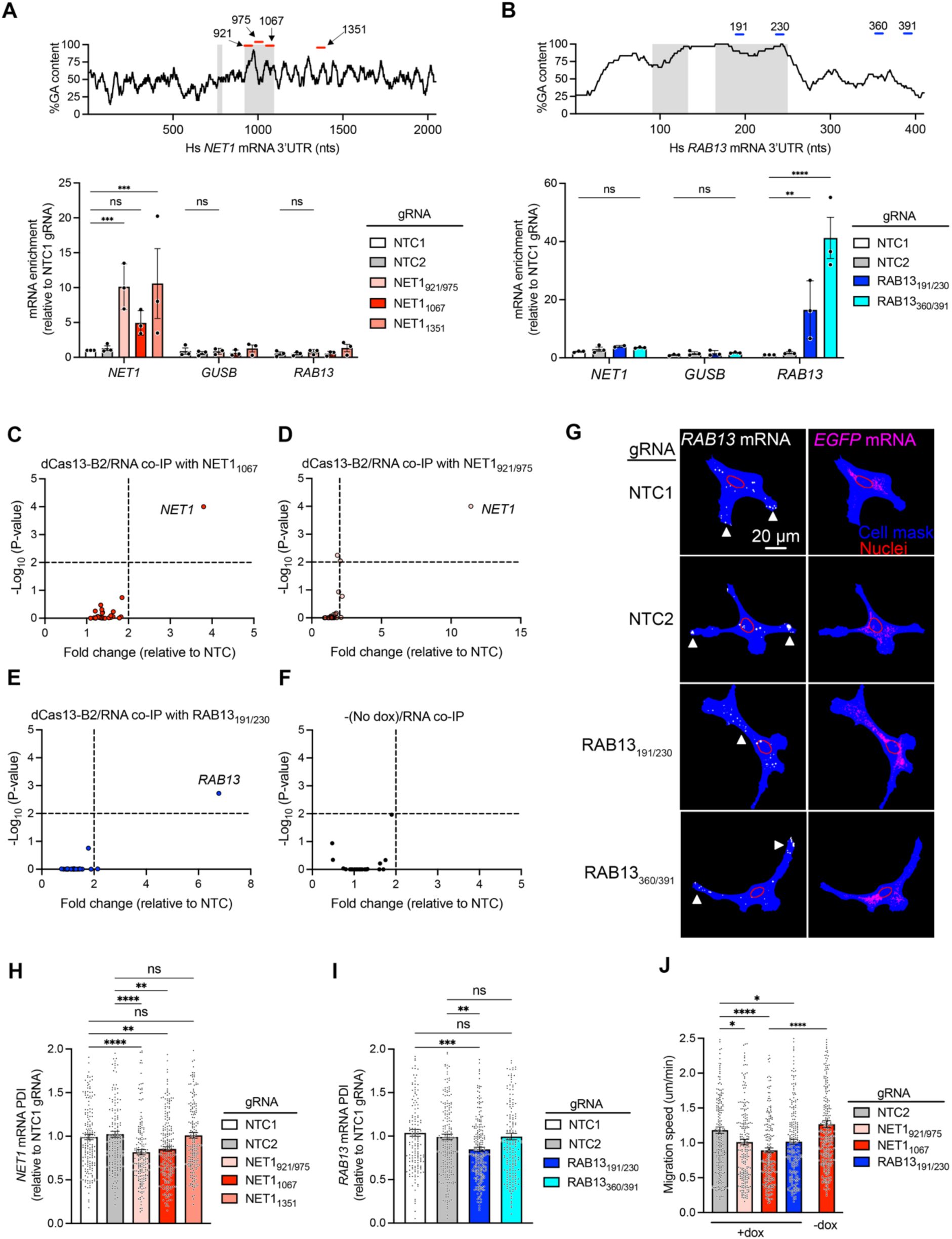
dCas13-B2 interferes with mRNA localization and mesenchymal cell migration in a sequence-specific manner. (**A**)-(**B**) (Top) GA-content in 15 nt increments along the *NET1* (A) and *RAB13* (B) 3’UTRs. Gray shading indicates previously described regions important for transcript localization. Red lines: gRNA target sites. (Bottom) Enrichment of *NET1, GUSB* and *RAB13* mRNAs, measured by ddPCR, in dCas13-B2 IPs with the indicated gRNAs. n = 3 independent experiments per condition. ** *P* < 0.001, *** *P* < 0.002, and **** *P* < 0.0001 by two-way ANOVA with Dunnett’s multiple comparison test. Representative blot of IP is found in Supplementary Figure S2. (**C**)-(**F**) Volcano plot of mRNA enrichments, by nanostring nCounter, in dCas13-B2 IPs with the indicated gRNAs. See Supplementary Table S3 for a full list of detected mRNAs. n = 3 independent experiments. Statistical comparisons were made relative to the NTC2 gRNA by two-way ANOVA followed by Dunnett’s multiple comparison test. (**G**) Representative FISH images of *RAB13* and *GFP* mRNA. Arrowheads indicate areas of *RAB13* mRNA accumulation. (**H**) and (**I**) PDI of *NET1* (H) and *RAB13* mRNA (I) in dCas13-B2 expressing cells with the indicated gRNAs. n = 174-223 (H) and n = 157-225 (I) cells per condition from 5-6 independent experiments; ** *P* < 0.01, *** *P* < 0.001, and **** *P* < 0.0001 by Kruskal-Wallis one-way ANOVA with Dunn’s multiple comparisons test. (**J**) Average migration speed of dCas13-B2 (+dox) expressing cells transfected with the indicated gRNAs. n = 212-287 cells per experimental condition. * *P* < 0.05, *** *P* < 0.005, and **** *P* < 0.0001 by Brown-Forsythe and Welch ANOVA multiple comparisons test. Bar graphs in panel (A), (B), and (H)-(J) represent mean ± SEM with an overlay of individual data points.

The ability of these gRNAs to target dCas13-B2 to specific mRNAs was assessed in co-immunoprecipitation experiments followed by ddPCR detection (Supplementary Fig. S2). Indeed, all NET1 gRNAs led to a significant enrichment of *NET1* mRNA in dCas13-B2 IPs, while non-targeted transcripts (*GUSB, RAB13*) were not appreciably enriched (Fig. 2A). In contrast, RAB13 gRNAs led to a significant enrichment of *RAB13*, but not of *NET1* or *GUSB* mRNAs (Fig. 2B). Non-specific interactions between dCas13 and non-target mRNAs have been described previously and are often, but not exclusively, attributable to gRNA sequence [37]. To further assess the targeting specificity of dCas13-B2 we designed nanoString nCounter panels that included: a) potential off-target transcripts of each gRNA based on partial complementarity; b) previously described Cas13d off-targets [37]; and c) additional mRNAs with GA-rich regions, like *NET1* and *RAB13*, which are also trafficked to cell protrusions by KIF1C (see Supplementary Table S3 for a full list). Importantly, the only mRNAs substantially enriched in dCas13-B2 co-immunoprecipitation samples were the targets of each gRNA (Fig. 2C-E; significance threshold > 2-fold enrichment and p ≤ 0.01; Source data are in Supplementary Table S3) while all other RNAs were detected only at background levels, similar to those seen in the absence of dCas13-B2 induction (Fig. 2F). Therefore, dCas13-B2 exhibits minimal off-target binding.

To determine whether dCas13-B2 binding to *NET1* and *RAB13* UTRs affects trafficking of these mRNAs, we used FISH to visualize and quantify the corresponding mRNA distributions (Fig. 2G-I). While as shown above, all dCas13-B2/gRNAs can bind either *NET1* or *RAB13* mRNA, only those gRNAs targeting the functionally important GA-rich regions (NET1_921/975_, NET_1067_ and RAB13_191/230_) led to a significant decrease in PDI indicating impairment of mRNA trafficking (Fig. 2H and I). A consequence of reduced *NET1* and *RAB13* peripheral localization is perturbation of cell migration and reduction in the speed of cellular movement [12,13,27]. Consistent with this, the decreased RNA trafficking upon dCas13-B2 binding to *NET1* and *RAB13* GA-rich regions was accompanied by reduced speed of random cell migration (Fig. 2J). The observed reduction was comparable to that reported previously using ASOs to disrupt trafficking of the respective mRNAs (Supplementary Fig. S3). Therefore, dCas13-B2 binds in a highly sequence-specific manner and affects the function of RNA elements to a degree sufficient to alter cellular behaviors.

### dCas13-B2 targeting to a GA-rich element interferes with CNBP recruitment

Given that dCas13-B2 can stably bind and functionally alter mRNA trafficking we hypothesized that dCas13-B2/gRNA complexes work by competing with RBP binding to GA-rich elements (Fig. 1B). To directly test this prediction, we turned to an in vitro system using purified components to assess whether dCas13-B2 competes with CNBP binding to target transcripts (Fig. 1B). In this system, an in vitro transcribed RNA fragment corresponding to the *RAB13* 3’UTR was designed to contain the λ bacteriophage BoxB hairpin in its 3’ end. The BoxB sequence is recognized by a peptide of the λN protein which when fused to GST allows for pulldown of the RNA and any associated factors on glutathione beads (Fig 3A). First, we confirmed that bacterially expressed and purified CNBP bound to the RAB13 3’ UTR with an apparent Kd of 0.9 uM (Supplementary Fig. S4A and B). Deletion of the GA-rich region in the RNA reduced CNBP binding (Supplementary Fig. S4A). These results are consistent with the previously reported direct interaction of CNBP with the *RAB13* RNA in vivo, and with the role of the GA-rich region in promoting CNBP binding [26]. Additionally, the observed Kd is within range of what has been previously reported for CNBP using other approaches and RNA substrates [32,38].

**Figure 3.**
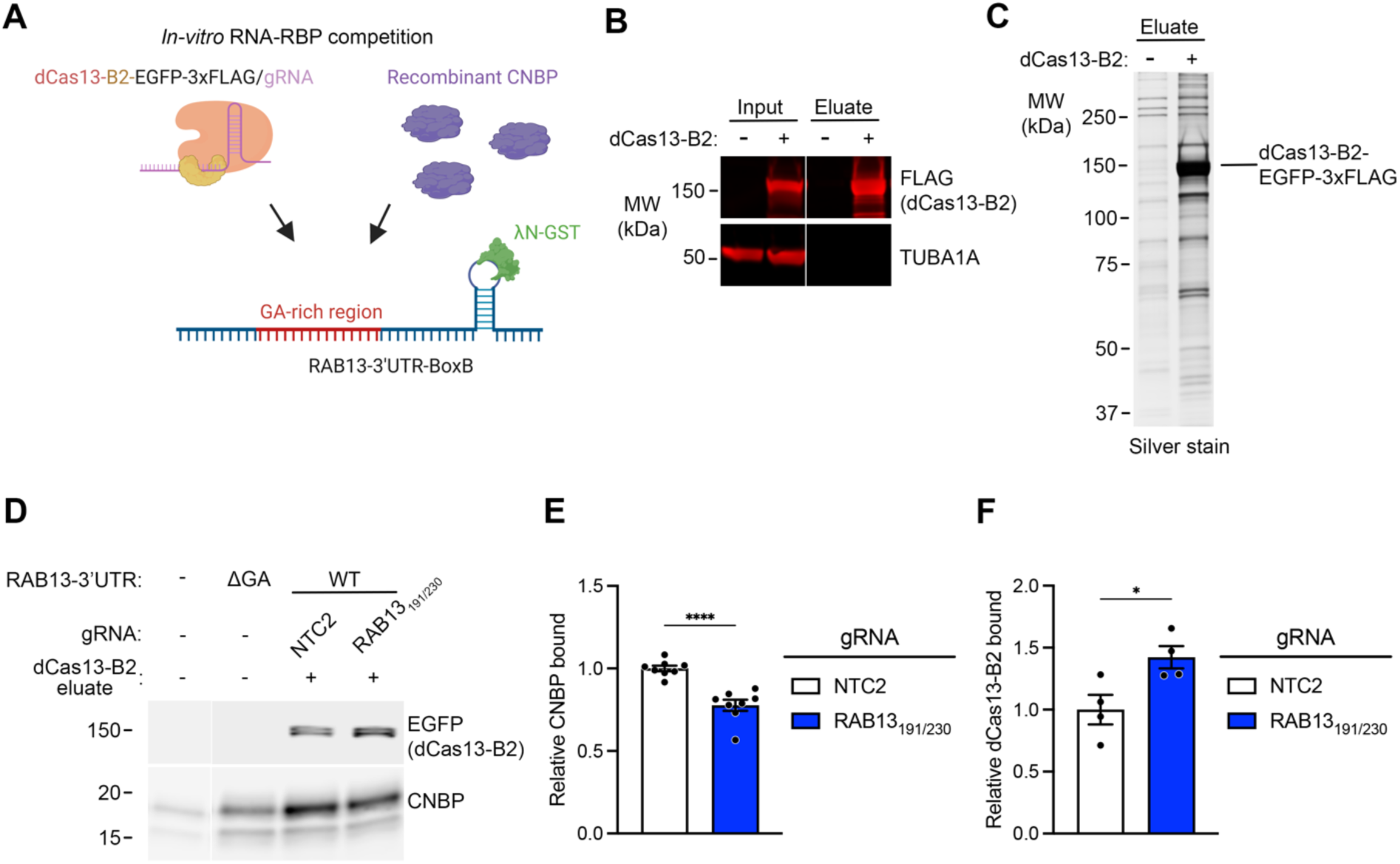
dCas13-B2/gRNA targeting to GA-rich regions competes with CNBP binding. (**A**) Schematic of the *in vitro* RNA-RBP binding assay. A box B stem-loop sequence downstream of the *RAB13* 3’UTR is used to pull down the RNA using λN-GST and assess competitive binding of dCas13-B2/gRNA and recombinant CNBP. (**B**) Representative western blot showing enrichment of dCas13-B2 upon immunoprecipitation with FLAG beads followed by elution of the protein with FLAG peptide. (**C**) Silver stain gel image of eluate with the apparent molecular weight of dCas13-B2 indicated to the right. (**D**) Representative western blot of bound GFP (dCas13-B2) and CNBP upon *RAB13* RNA pulldown. Note that dCas13-B2/gRNA complex was only added to the WT RNA samples. Also, binding in the presence of NTC2 gRNA stems largely from background association with the beads (see Supplementary Fig. S4D). (**E**)-(**F**) Quantification of bound CNBP and dCas13-B2 normalized to WT *RAB13* 3’UTR as in (D). n = 8 (E; CNBP) and n = 4 (F; dCas13-B2) independent experiments; * *P* < 0.05 and *** *P* < 0.0001 by two-tailed Student’s unpaired t-test. Bar graphs in panel (E)-(F) represent mean ± SEM with an overlay of individual data points.

To isolate dCas13-B2/gRNA complexes, the protein was produced by transient transfection in HEK293 cells, immunoprecipitated on anti-FLAG antibody beads, and loaded with synthetic gRNA. Free gRNA was removed and dCas13-B2/gRNA complexes were eluted using excess FLAG peptide leading to their significant enrichment (Fig. 3B and C; Supplementary Fig. S4C). We then assessed how dCas13-B2 affected the binding of CNBP to the RAB13 3’UTR. We note that the maximal amount of dCas13 that we could add in the reaction was sub-stoichiometric to CNBP (Supplementary Fig. S4C). Nevertheless, despite a limiting quantity, dCas13/RAB13_191/230_ complexes significantly reduced association of CNBP with the target RNA (Fig. 3D and E), while at the same time increasing association between the RNA and dCas13-B2 (Fig. 3D, F and Supplementary Fig. S4D). These results support the notion that dCas13-B2/gRNA complexes can compete with RBP recruitment on an RNA target. We suggest that dCas13-B2 can be used as an effective, specific and versatile tool to manipulate specific RNA-RBP interactions and investigate the functional contributions of RNA elements.

### Nuclear retention and expression levels limit the efficacy of UC-driven gRNAs

Our characterization thus far had relied on transfection of chemically synthesized gRNAs into cells. However, this approach has the same limitations presented by ASO delivery (i.e. inability to assess long term effects in dividing cells or tissues, and promiscuous uptake; see Introduction). For CRISPR-dCas13 to provide an improved ASO alternative, all required components (i.e., dCas13 as well as the gRNA) should be genetically encoded and able to be expressed in a stable manner. We, therefore, used lentiviral transduction to stably integrate a gRNA expression cassette (under a human U6 promoter; U6-gRNA), in cells also expressing dox-inducible dCas13-B2. Dox induction was initiated 48 hrs prior to assay. However, in contrast to transfected gRNA we noticed that U6-driven NET1_1067_ gRNA did not have a detectable effect on *NET1* mRNA localization (Supplementary Fig. S5A), which remained similar to that of control cells (expressing U6-NTC1 gRNA). No effect was detected even upon increasing the multiplicity of infection (MOI) 10 times (Supplementary Fig. S5A).

We reasoned that the ability of a gRNA to direct dCas13-B2 binding to a target RNA and compete with RBP recruitment likely relies on the amount of gRNA. Additionally, given that the RNA localization regulation we study occurs in the cytoplasm, gRNA presence in the cytoplasm could also be a limiting factor. We thus used ddPCR to assess gRNA amount and the RNAscope Plus smRNA assay to visualize gRNA distribution. Cell lines expressing U6-driven gRNAs, either NTC1 or NET1_1067_, exhibited gRNA amounts that were orders of magnitude lower compared to the amount detected upon synthetic gRNA transfection (Supplementary Fig. S5B and C). Higher MOI led to only a slight increase, which plateaued rapidly (Supplementary Fig. S5B and C). To visualize gRNA in situ we designed smRNA probes against the U6-NET1_1067_ gRNA. Specific signal was readily detectable in stably expressing cells but absent from the parental cell population (Supplementary Fig. S5D). Interestingly, though, the signal was detected primarily in the nucleus, indicating that the expressed gRNA was likely not efficiently exported to the cytoplasm (Supplementary Fig. S5D). Overall, it appeared that both insufficient cytoplasmic accumulation and low expression levels of U6-driven gRNAs limit the ability of CRISPR-dCas13 to functionally interfere with cytoplasmic RBP-RNA interactions.

To increase gRNA levels, we expressed a gRNA array in which a U6 promoter drives transcription of a precursor containing 8-10 gRNA sequences separated by direct repeats. Individual gRNAs can be cleaved from such arrays through the catalytic action of Cas13 (Fig. 4A). Importantly, the dCas13 variant, which does not cleave mRNA due to mutations in the HEPN domain, still retains gRNA array processing activity [29]. Consistent with this, Northern blot analysis showed that dCas13-B2 was able to process U6-gRNA arrays into mature gRNAs (Fig. 4B).

**Figure 4.**
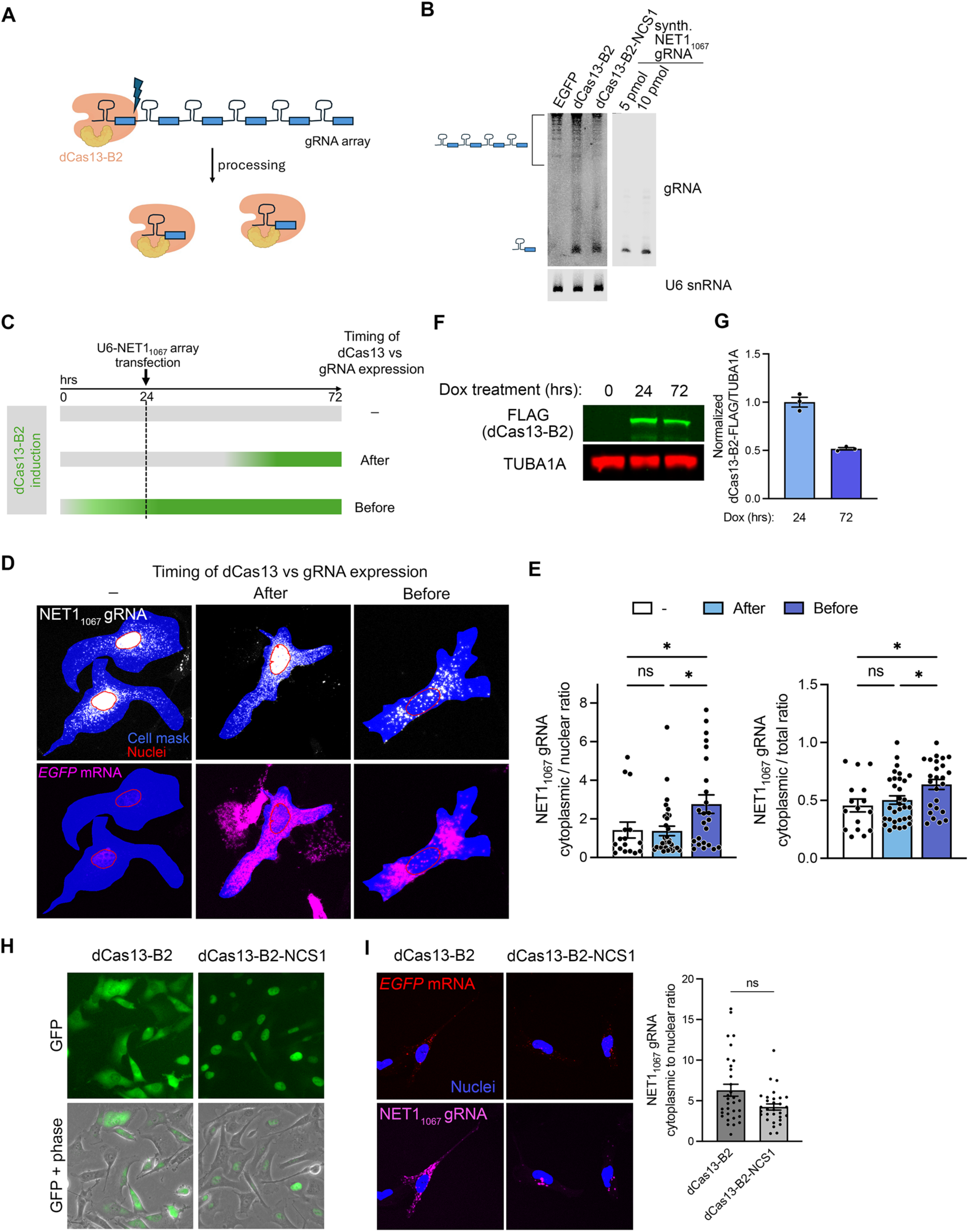
Prolonged dCas13-B2 expression is required for cytosolic enrichment of gRNAs processed from multimeric arrays. (**A**) Schematic of gRNA array processing by dCas13-B2. (**B**) Representative Northern blot detection of NET1_1067_ gRNA from total RNA of HEK293 cells expressing the U6-NET1_1067_ gRNA array and the indicated proteins. A fragment comparable in size to a synthetic gRNA was detectable in cells expressing dCas13-B2 constructs, indicating processing of the array. U6 snRNA served as a loading control. (**C**) Schematic of experimental set up to test the effect of timing of dCas13-B2 expression for gRNA accumulation in the cytosol. MDA-MB-231 cells used were stably integrated with Dox-inducible dCas13-B2. (**D**) Representative FISH images of NET1_1067_ gRNA and *GFP* mRNA detected by RNAscope, under conditions described in (C). Area of cells expressing both *GFP* mRNA and gRNA is shown in blue. Nuclear outlines are shown in red. (**E**) Amount of cytoplasmic NET1_1067_ gRNA relative to nuclear (left) or entire cell amount (right). n = 16-31 cells per condition as described in (C); * *P* < 0.05 by one-way ANOVA with Holm-Sidak’s multiple comparison test. (**F**)-(**G**) Representative western blot and quantification showing that dCas13-B2 expression peaks 24 hours after induction. n=3 independent experiments. (**H**) Representative GFP fluorescence and phase images of dCas13-B2 and dCas13-B2-NCS1 expressing cells showing that NCS1 fusions are predominantly nuclear. (**I**) MDA-MB-231 cells were transiently transfected with the indicated dCas13-B2 constructs and the U6-NET1_1067_ gRNA array. Representative FISH images show NET1_1067_ gRNA and *GFP* mRNA with quantification of cytoplasmic to nuclear gRNA amount. n = 30-32 cells per condition; not significant (ns) by one-way ANOVA with Kruskal-Wallis multiple comparisons test. Bar graphs in panel (G), (E), and (I) represent mean ± SEM with an overlay of individual data points.

Nevertheless, in situ gRNA detection showed that a significant amount of gRNA was still retained in the nucleus. Specifically, when the U6-gRNA array cassette was delivered into cells and then expression of dCas13-B2 was induced (Fig. 4C, ‘After’), almost half of the gRNA signal was present in the nucleus (Fig 4D and E). Intriguingly, this gRNA distribution was similar to when dCas13-B2 was not induced at all (Fig. 4D and E). We reasoned that dCas13-B2 expression prior to gRNA array expression would be required to alter gRNA distribution. To test that, dCas13 expression was induced prior to transfection of the U6-gRNA array cassette (Fig. 4C, ‘Before’). Interestingly, this led to a substantial increase in cytoplasmic gRNA together with a decrease in nuclear gRNA (Fig. 4D and E). The effect was not due to higher dCas13-B2 expression, as dCas13-B2 expression peaks at 24 hrs (Fig. 4F and G). While we cannot conclude on the exact mechanistic basis for the increased cytoplasmic presence of gRNA, these data suggested that the timing or duration of dCas13-B2 expression matters. Short-term expression of dCas13 is not optimal for achieving higher amount of cytoplasmic gRNA, an element which is required for interfering with RBP-RNA interactions that occur in the cytoplasm.

We further investigated whether increasing the presence of dCas13-B2 in the nuclear compartment would affect gRNA distribution [39]. For this, the protein was additionally fused to a nucleocytoplasmic shuttling (NCS1) module, consisting of two SV40 nuclear localization signals and a leucine-rich nuclear export signal [39]. As expected, the dCas13-B2-NCS1 protein exhibited a more pronounced nuclear accumulation (Fig. 4H) and was also able to process gRNA arrays (Fig. 4B). Nevertheless, it did not further improve the nucleocytoplasmic gRNA distribution (Fig. 4I). We thus carried on with the dCas13-B2 fusion without including additional trafficking signals.

### Stable expression of a genetically-encoded CRISPR-dCas13 system interferes with GA-element-dependent mRNA trafficking

Relying on the above information, we stably integrated gRNA arrays (containing NTC1 or NET1_1067_ gRNAs) either into cells constitutively expressing dCas13-B2 (CMV_dCas13-B2) or into cells with inducible dCas13-B2 (pIND20_dCas13-B2). In the latter case, dCas13-B2 expression was induced for 2 or 7 days prior to assay. The amount of gRNA expressed was similar in all cases and higher than that achieved with a single U6-gRNA cassette (Fig. 5A), confirming that U6-gRNA arrays enhance gRNA production. Inducible dCas13-B2 expression already plateaued by 2 days (Supplementary Fig. S6A) and was observed in only a fraction of the cells, in contrast to the CMV_dCas13-B2 expressing cells, which exhibited a more uniform expression (Supplementary Fig. S6B). Nevertheless, individual GFP-positive cells exhibited similar amounts of dCas13-B2-GFP expression (Supplementary Fig. S6C). Interestingly, CMV_dCas13-B2 cells exhibited significantly higher gRNA amount in the cytoplasm, compared to Dox-induced_dCas13-B2 cells (Fig. 5B and C). This difference was not simply due to overall differences in gRNA amount as normalization to total gRNA showed similar trends (Supplementary Fig. S6D). These results are consistent with the notion that short-term expression of dCas13 is not optimal for achieving higher amounts of cytoplasmic gRNA. Indeed, prolonged induction of dCas13-B2 for 7 days increased cytoplasmic gRNA levels, even though not to the same extent as in constitutive expressing cells (Fig. 5B and Supplementary Fig. 6D).

**Figure 5.**
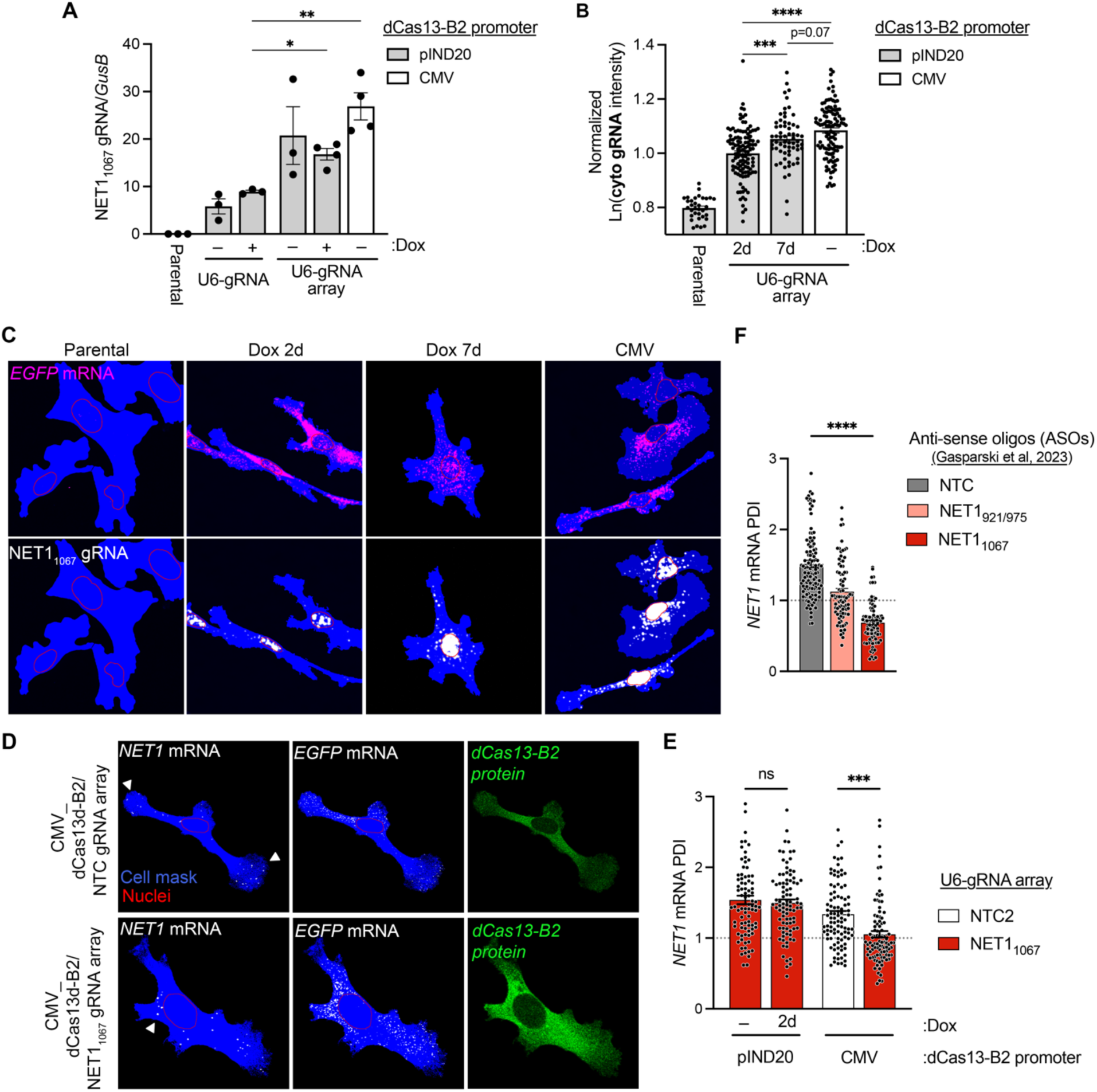
A stably integrated dCas13-B2/gRNA array system alters endogenous *NET1* mRNA distribution. (**A**) ddPCR quantification of NET1_1067_ gRNA normalized to the house keeping gene *GusB*. U6-gRNA array cell lines exhibit higher overall gRNA abundance. n = 3-4 independent experiments; ** *P* < 0.01 by one-way ANOVA with Holm-Sidak’s multiple comparisons test. (**B)** Cytoplasmic NET1_1067_ gRNA natural logarithm (ln) intensities (from FISH images as in (C) in stable cell lines with inducible (pIND20) or constitutive (CMV) dCas13-B2 expression. Days of Dox induction are indicated. n = 65-115 cells per condition from 2 independent experiments; *** *P* < 0.001 and **** *P* < 0.0001 by one-way ANOVA with Sidak’s multiple comparisons test. (**C**) Representative FISH images of NET1_1067_ gRNA and *GFP* mRNA, associated with (B). Cell area is shown in blue. Nuclear outlines are shown in red. (**D**) Representative images of stable cell lines constitutively expressing dCas13-B2 and the indicated gRNA arrays. *NET1* and *GFP* mRNAs are detected by FISH, and dCas13-B2 protein by GFP fluorescence. Areas of *NET1* mRNA accumulation are indicated by white arrowheads. (**E**) PDI of *NET1* mRNA across indicated stable cell lines. n = 86-99 cells per condition from 3-4 independent experiments; *P* < 0.0001 by one-way ANOVA with Sidak’s multiple comparisons test. (**F**) Data from Gasparski et al. (2023), showing effect of ASOs on *NET1* mRNA PDI, for comparison. n = 84-118 cells per condition from 3 independent experiments; *P* < 0.0001 by one-way ANOVA with Sidak’s multiple comparisons test. Bar graphs in panel (A), (B), (E) and (F) represent mean ± SEM with an overlay of individual data points.

To assess whether these differences in cytoplasmic gRNA amount influenced the effectiveness of dCas13-B2 to compete with GA-element-dependent mRNA targeting, we measured *NET1* mRNA localization. In accordance with cytoplasmic gRNA amounts, short-term dCas13-B2 induction, for 2 days, did not lead to a detectable effect on *NET1* mRNA localization (Fig. 5E), while constitutive expression led to a significant reduction in peripheral *NET1* mRNA localization (Fig. 5D and E). The extent of *NET1* mRNA mislocalization in NET1_1067_ gRNA expressing cells constitutively expressing dCas13-B2 was similar to that observed upon synthetic gRNA delivery (compare Fig. 5E with Supplementary Fig. S5A), and similar to previously reported ASO-mediated effects (Fig. 5F). Prolonged, 7-day induction led to a detectable, albeit not as robust effect (Supplementary Fig. S6 E-G).

### Preventing NET1 mRNA trafficking reduces cancer cell invasion and metastatic lung colonization

To ensure that this system does not activate cellular stress pathways we compared parental and dCas13-B2 lines. Notably there was no detectable difference in the stress response, as indicated by eIF2α phosphorylation under either basal conditions or after sodium arsenite treatment (Supplementary Fig. S7A). Further supporting this conclusion, RNA-seq analysis revealed that expression of genes associated with cellular stress and cytotoxicity was not appreciably altered in all modified cell populations (Supplementary Fig. S7B and Supplementary Table S4).

Being able to genetically encode all system components opened the possibility of interrogating the consequences of altering *NET1* mRNA localization in long-term phenotypic assays. Using the constitutively expressing CMV_dCas13-B2 cells, we found that perturbation of *NET1* mRNA localization did not affect the growth rate of cells in 2D culture conditions (Fig. 6A). However, embedding single cells in 3D matrigel and allowing them to form multicellular spheroids over ∼9 days (Fig. 6B) revealed that cells expressing the NET1_1067_-gRNA array formed smaller spheroids (Fig. 6C), indicating reduced growth or viability in these 3D growth conditions. To assess invasive potential, serum withdrawal was used to induce collective invasion in the surrounding matrix [28] (Fig. 6B). Consistent with a prior report using ASOs [28], altering *NET1* mRNA localization through dCas13-gRNA expression led to significantly reduced spheroid invasiveness (Fig. 6D).

**Figure 6.**
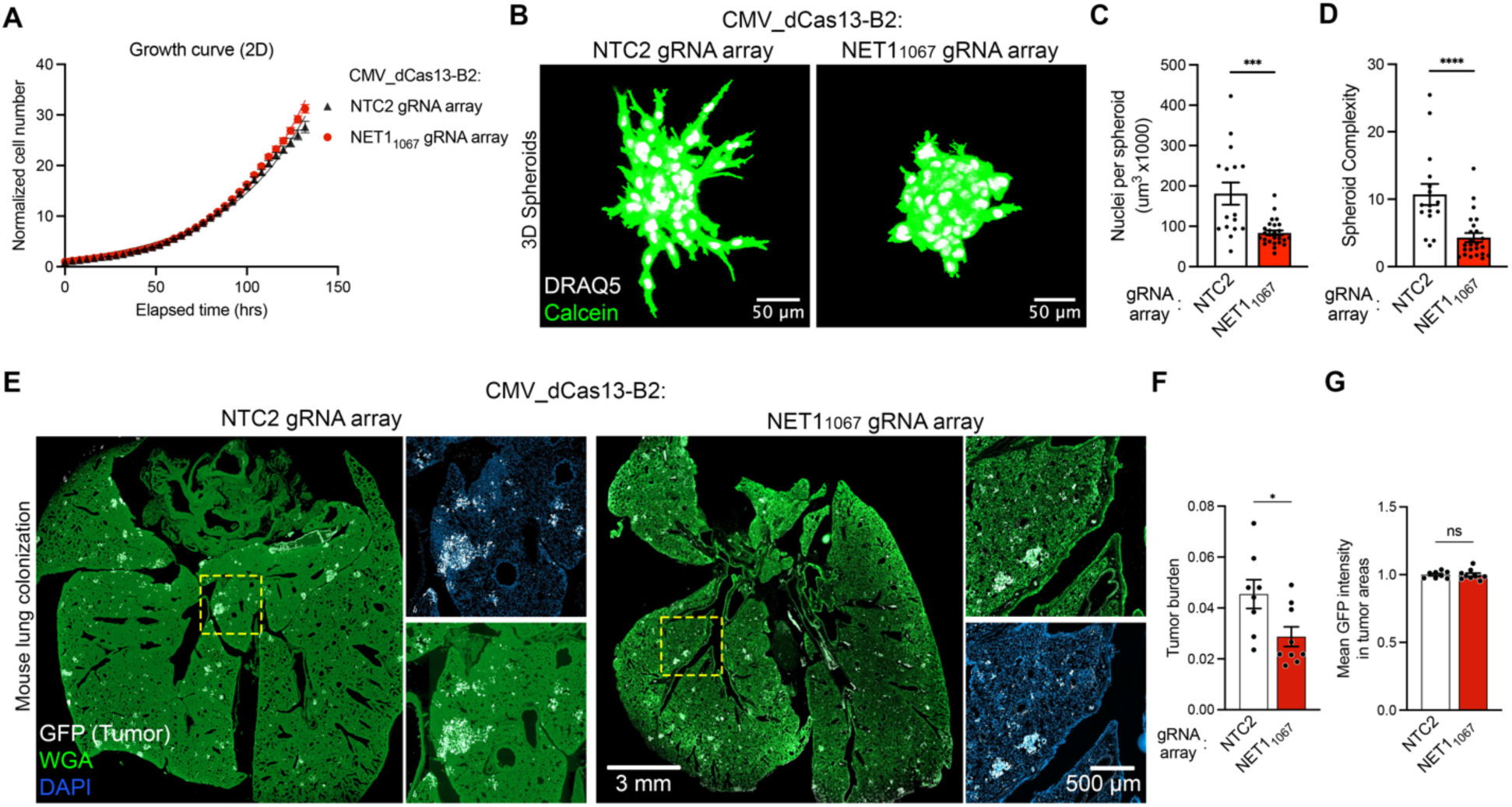
Stable *NET1* mRNA mislocalization leads to reduced 3D invasion and metastatic lung colonization. (**A**) Proliferation during 2D culture of CMV_dCas13-B2 cells stably expressing the indicated gRNA arrays, measured by Incucyte confluence analysis. (**B**) Single cells of the indicated stable lines were embedded in 3D Matrigel. Images show representative spheroids formed after 9-day growth and overnight serum withdrawal to induce collective invasion. Cells were stained with Calcein-AM dye, nuclei with Draq5. (**C**) Total nuclear volume per spheroid, indicating the number of cells per spheroid. (**D**) Spheroid complexity, defined as Perimeter^2^/(4π x Area), indicating degree of invasiveness. n = 16 or 26 spheroids per condition from 2 independent experiments; *** *P* < 0.001 and **** *P* < 0.0001 by Mann-Whitney test. (**E**) Representative images of lung tissue sections from mice intravenously injected with the indicated cells through the tail vein. Lung tissue was stained with WGA. GFP immunofluorescence was used to identify tumor colonies. (**F**) Tumor burden, calculated as the ratio of GFP-positive over normal tissue area. (**G**) Mean GFP intensity in tumor areas per animal. n = 8-9 mice per condition. * *P* < 0.05 by two-tailed Student’s unpaired t-test.

To further extend these observations and assess a potential role of *NET1* mRNA localization in cancer metastasis in vivo, CMV_dCas13-B2-GFP cells were injected intravenously in the tail vein of immunodeficient mice. Metastatic colonization in the lungs was assessed 5 weeks later, through immunofluorescent detection of GFP (Fig. 6E). While dCas13-B2-GFP protein levels in tumor areas remained similar, expression of the NET1_1067_-gRNA array led to significantly reduced tumor burden (Fig. 6F, G).

Overall, the data described here demonstrate that dCas13 can be used to effectively perturb specific RNA-RBP interactions and assess functional contributions. We additionally show that the effective implementation of this CRISPR-dCas13 tool in a stably-expressed, fully genetically encoded manner relies on certain considerations. While some CRIPSR-Cas13 applications can be achieved through transient binding with target mRNAs, the persistent interference with endogenous RBPs requires stable interaction, which can be enhanced by fusion to dsRBDs [21]. We find the B2 protein dsRBD to provide maximal effect in this context. An additional consideration for targeting cytoplasmic mRNAs involves gRNA export from the nucleus and accumulation to sufficient levels in the cytoplasm. In contrast to transient expression, stable genomic integration of U6-driven gRNA constructs leads to cytoplasmic gRNA levels which can be limiting. We show that their levels can be further influenced by the constitutive or inducible nature and duration of dCas13 expression. Additional modifications including gRNA expression from a U1 small nuclear RNA promoter could further improve outcomes [40,41].

We envision that this dCas13-based tool can be used to broadly interrogate specific RNA-RBP binding events and can thus provide a flexible and reversible approach to delineate regulatory RBP contributions and dissect RNP organization without alteration of the genomic DNA. Nevertheless, specific optimizations and considerations might be required depending on the particular binding event under study. We have applied this approach to the study of mRNA localization driven by GA-rich elements. This targeting mechanism operates in a variety of cell types in in vitro cultures or in vivo tissues and is important for diverse processes ranging from cancer cell invasion to blood vessel patterning and epithelial tissue organization [23,24,28,42]. Here, we extend these findings to show that mistargeting of the GA-containing *NET1* mRNA leads to reduced metastatic lung colonization in mouse models. The described use of CRISPR-dCas13 to reprogram the distribution of specific mRNAs, as well as alternative methods to mis-target mRNAs to cytoplasmic locations [19], will allow further investigation into the long-term phenotypic consequences of this widespread mRNA localization pathway. By enabling durable perturbation of defined RNA regulatory elements, the platform described here opens avenues for mechanistic studies of post-transcriptional regulation in physiologically relevant and disease-associated contexts.

## DATA AVAILABILITY

All data supporting the findings of this study are available from the corresponding author on reasonable request.

## ACKNOWLEDGMENTS

We thank the CCR Genomics Core of the National Cancer institute, NIH, for ddPCR and nanoString nCounter analysis, and Yi He from the Protein Expression facility, NHLBI, NIH, for help with protein purification. Analysis and management of tissue images were supported by the NCI HALO Image Analysis Resource. The graphical abstract was originally created in BioRender, Mason D. (2026) https://BioRender.com/kdmxt2k and modified using ChatGPT (OpenAI). The authors reviewed and approved the resulting figure.

## AUTHOR CONTRIBUTIONS

D.E.M. and S.M. conceived the project. D.E.M., D.B., N.J. designed experiments. D.E.M., D.B., N.J., I.G.J. and B.M. performed experiments. D.E.M., D.B., N.J., R.I.-B. and S.M. analyzed data. D.E.M., D.B. and S.M. wrote original manuscript. All authors reviewed and edited the manuscript.

---

**Supplementary Figure S1.**
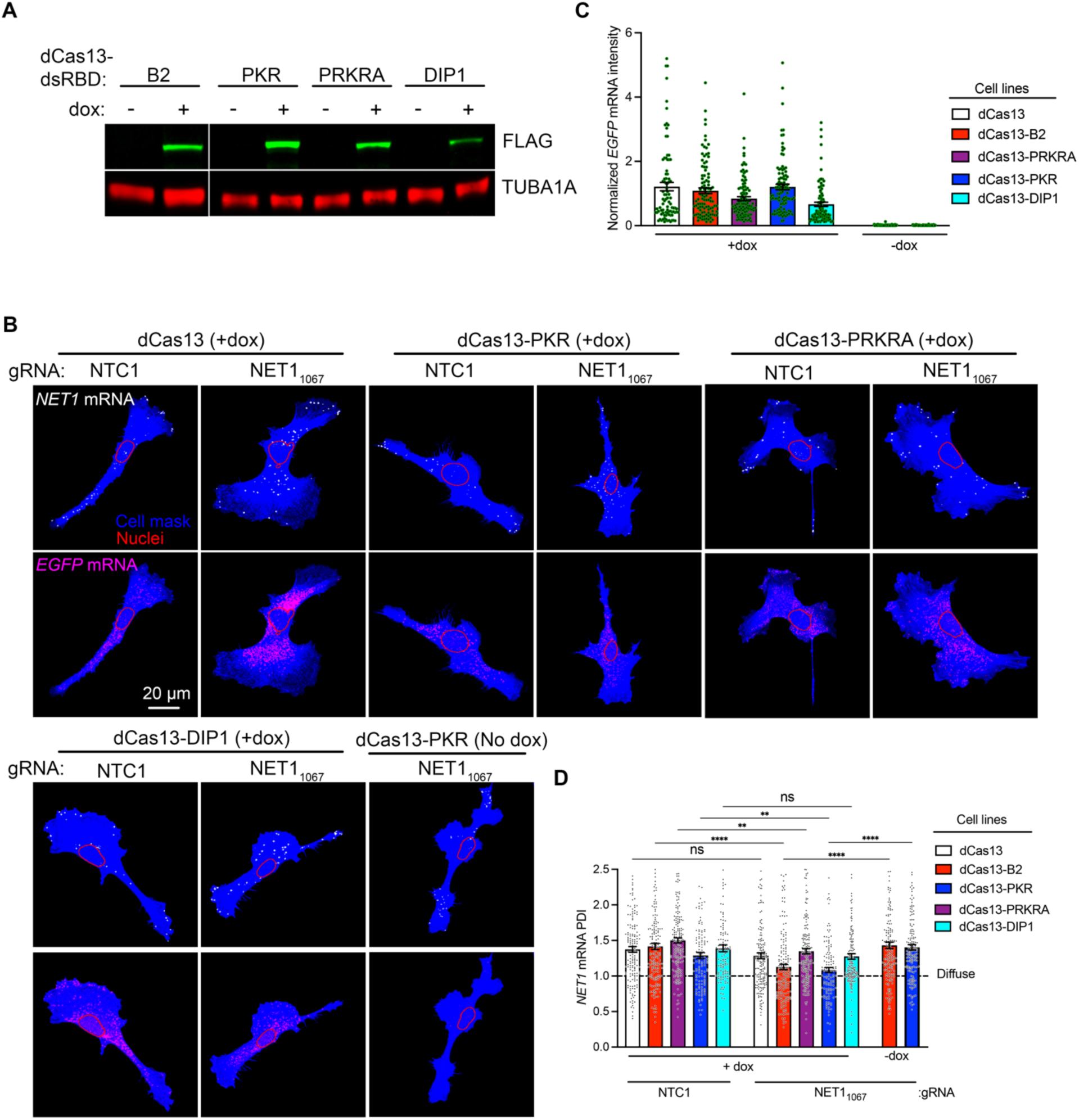
Fusion of dsRBDs alters the effect of dCas13/gRNA on *NET1* mRNA localization to cell protrusions. (**A**) Representative Western blot of dCas13-dsRBD expression (detected through FLAG tag) in Dox-inducible cell lines. (**B**) Representative FISH images of *NET1* and *GFP* mRNA in the indicated dCas13-dsRBD cell lines transfected with synthetic NTC1 or NET1_1067_ gRNAs. (**C**) Normalized *GFP* mRNA intensity from cells as in (B). n = 88-117 cells per condition. (**D**) PDI of NET1 mRNA in dCas13-dsRBD expressing cell lines (as in (B)). Note that PDI=1 reflects a diffuse distribution. n = 113-181 cells per condition from 4-6 independent experiments; ** *P* < 0.01 and **** *P* < 0.0001 by Kruskal-Wallis one-way ANOVA with Dunn’s multiple comparisons test.

**Supplementary Figure S2.**
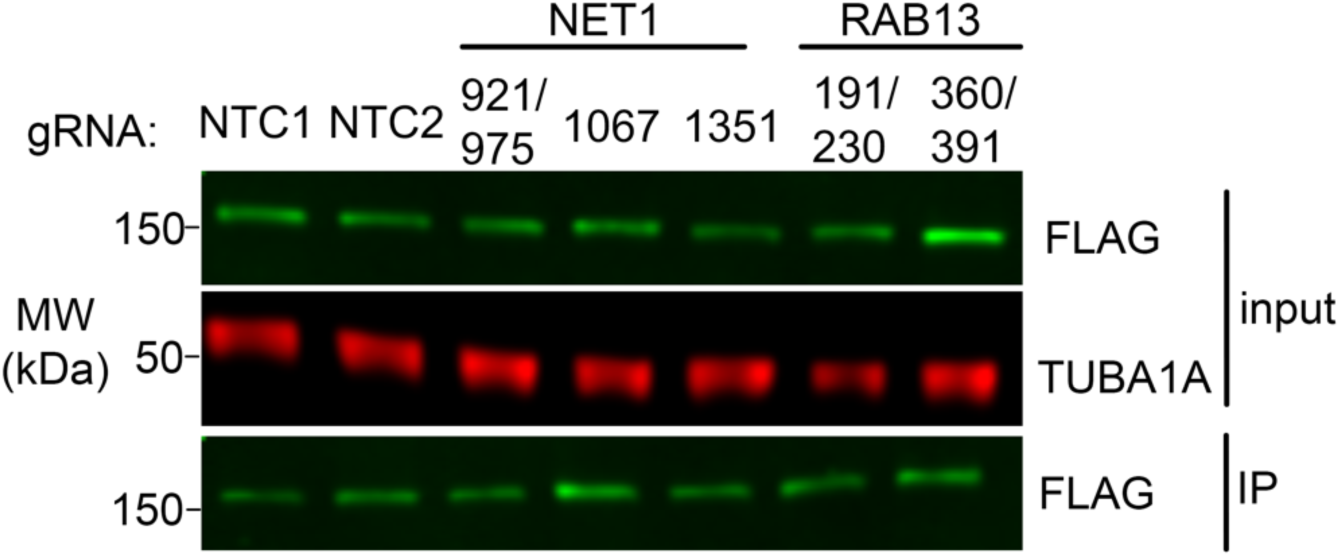
dCas13-B2 enrichment by immunoprecipitation. Representative Western blot of immunoprecipitated dCas13-B2, using FLAG-tag antibody, from cells also transfected with the indicated synthetic gRNAs (Related to Fig. 2 A and B).

**Supplementary Figure S3.**
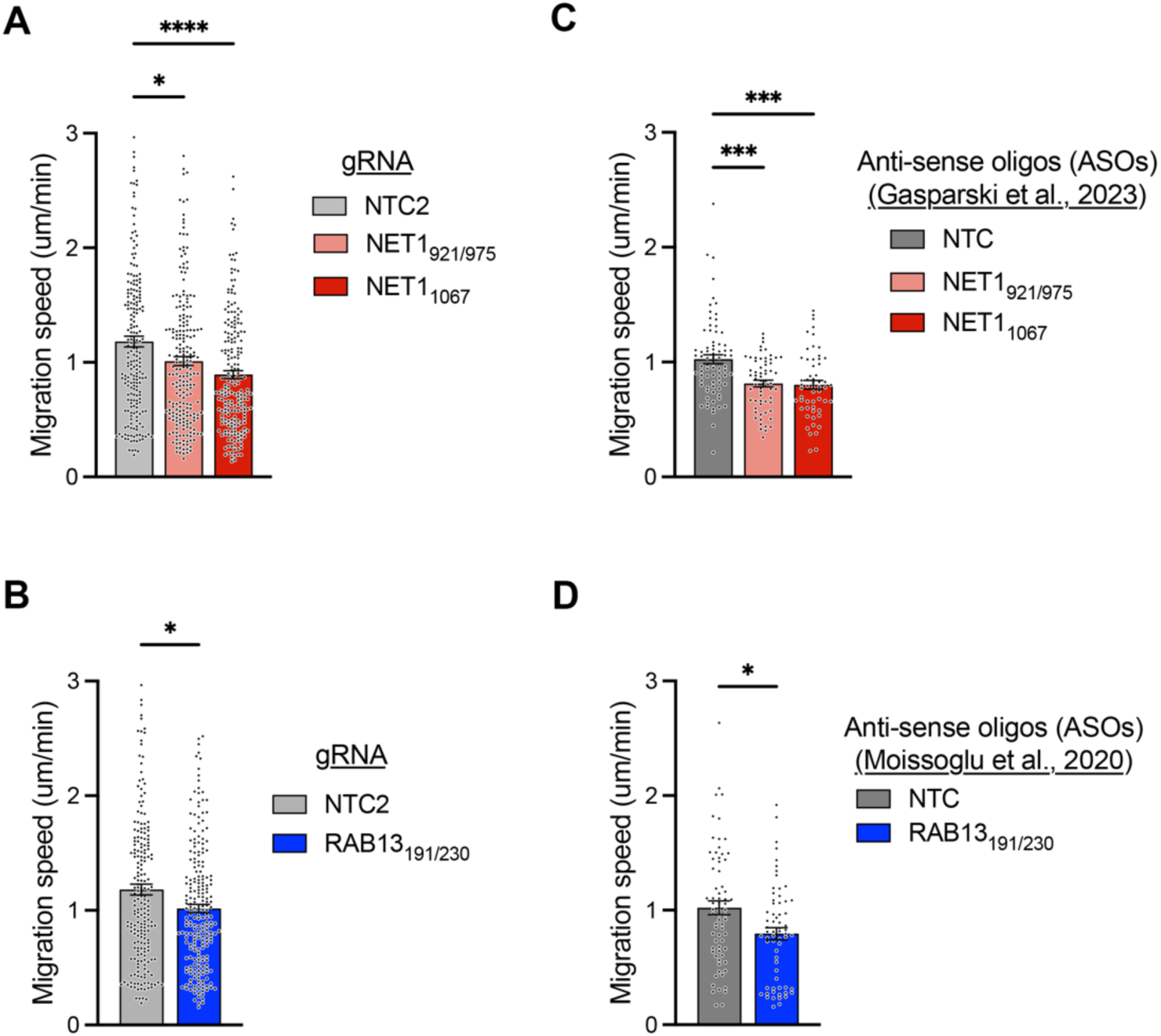
Effect of dCas13-B2/gRNA is comparable to that achieved upon antisense oligo delivery. (**A, B**) Data from Figure 2J. Average migration speed of dCas13-B2 expressing cells transfected with the indicated gRNAs against NET1 (A) or RAB13 (B). (**C, D**) Data from Gasparski et al. (2023). Average migration speed of MDA-MB-231 cells upon delivery of the indicated ASOs against NET1 (C) or RAB13 (D).

**Supplementary Figure S4.**
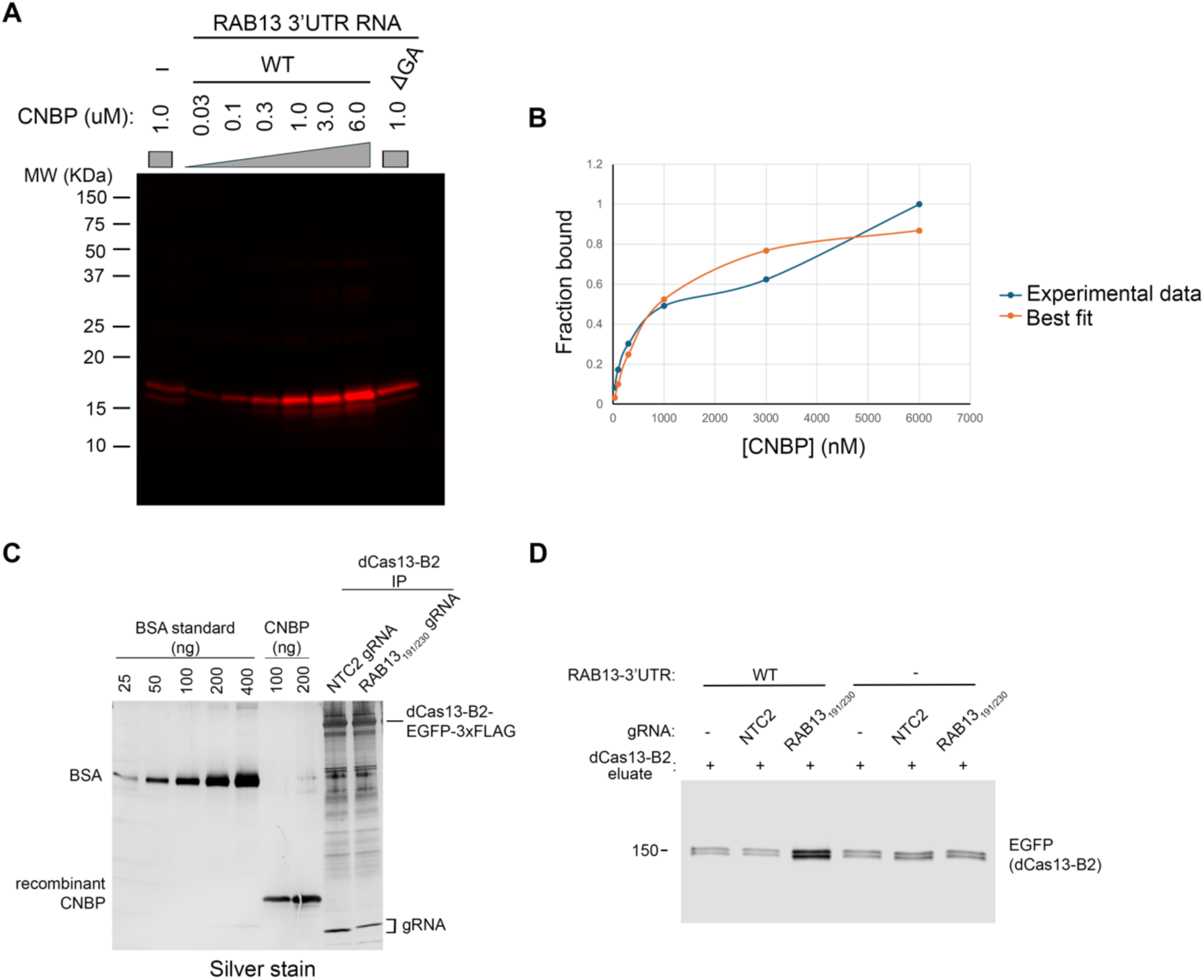
Binding of purified recombinant CNBP to the RAB13 3’UTR. (**A**) Representative western blot of recombinant CNBP bound to in vitro transcribed wild type (WT) or (DGA) RAB13-BoxB 3’UTR fragment. Bound protein was recovered after λN-GST pulldown. Increasing concentration of recombinant CNBP was used to calculate the apparent binding constant K_d_. (**B**) Quantification of CNBP bound to the WT RAB13 3’UTR fragment, from data as in (A). Apparent K_d_ is ∼0.9 μM, with the assumption that CNBP binds in a non-cooperative manner at a single-site on the Rab13 3’ UTR. (**C**) Silver stain gel image of purified CNBP and immunoprecipated dCas13-B2 loaded with the indicated gRNAs. Amounts are equivalent to those used in binding reactions shown in Fig 3D-F. Note that dCas13-B2 is sub-stoichiometric to CNBP. Bovine serum albumin is used as a reference. Silver-stained species likely corresponding to gRNAs are indicated. (**D**) Western blot of bound GFP (dCas13-B2) upon RNA pulldown. Note that there is detectable binding of dCas13-B2 even in the absence of target RNA (- *RAB13* 3’UTR) and/or gRNA, likely reflecting non-specific binding to the beads. Results are representative of two independent replicates.

**Supplementary Figure S5.**
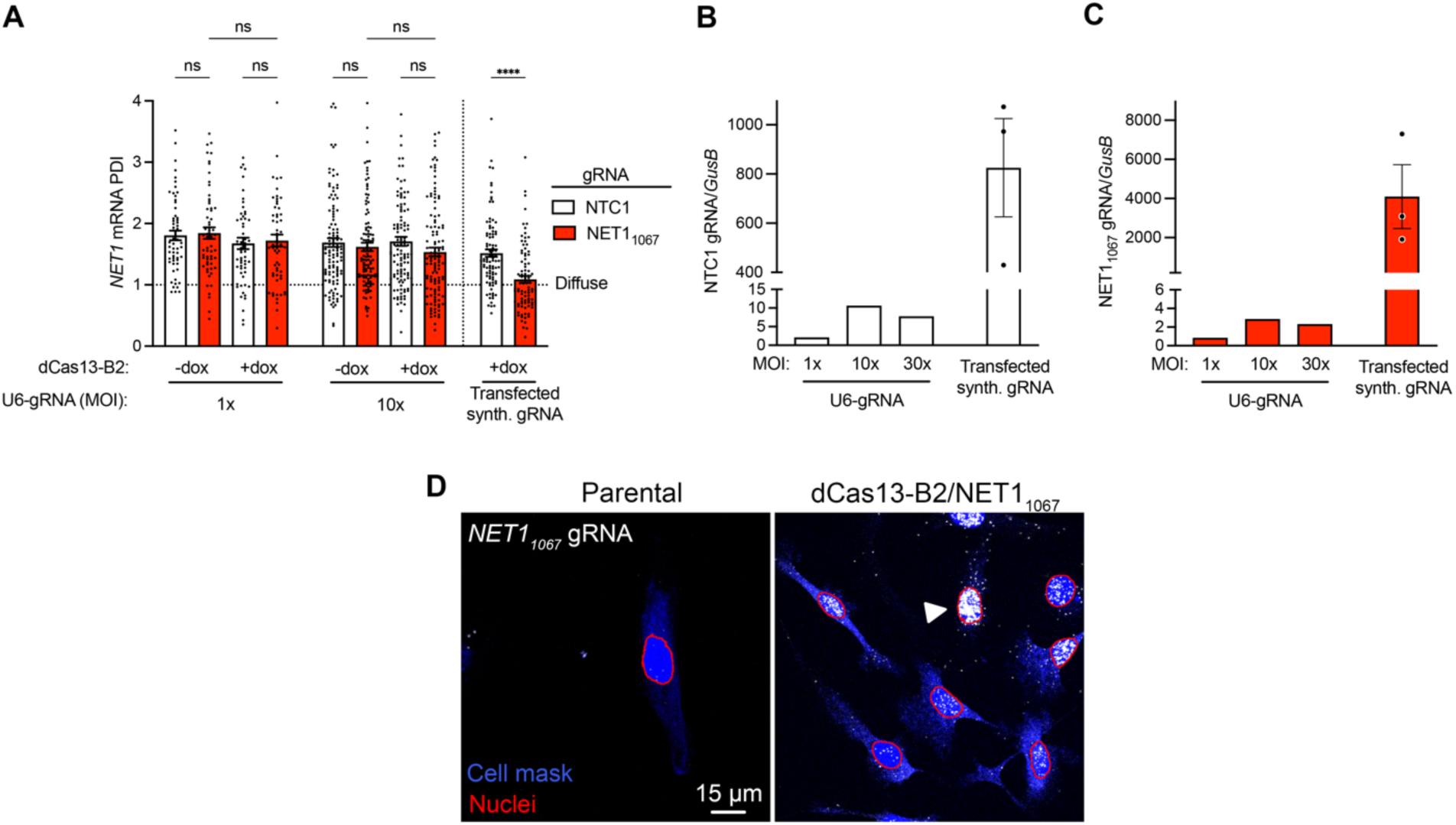
Stable integration of U6-gRNA cassette is insufficient to alter RNA localization, due to low expression and nuclear gRNA accumulation. (**A**) PDI of *NET1* mRNA in cell lines expressing single gRNA (NTC1 or NET1_1067_) from a U6 promoter. Effect of transfected synthetic gRNA is shown for comparison. n = 60-119 cells per condition; **** *P* < 0.0001 by Kruskal-Wallis ANOVA with Dunnett’s multiple comparisons test. (**B**)-(**C**) gRNA expression, by ddPCR, normalized to *GusB* mRNA for both NTC1 (B) and NET1_1067_ (C) gRNAs. (**D**) Representative FISH images of NET1_1067_ gRNA in indicated cell lines. Cell area shown in blue and nuclear outline in red. Arrowhead indicates accumulation of NET1_1067_ gRNA in the nucleus.

**Supplementary Figure S6.**
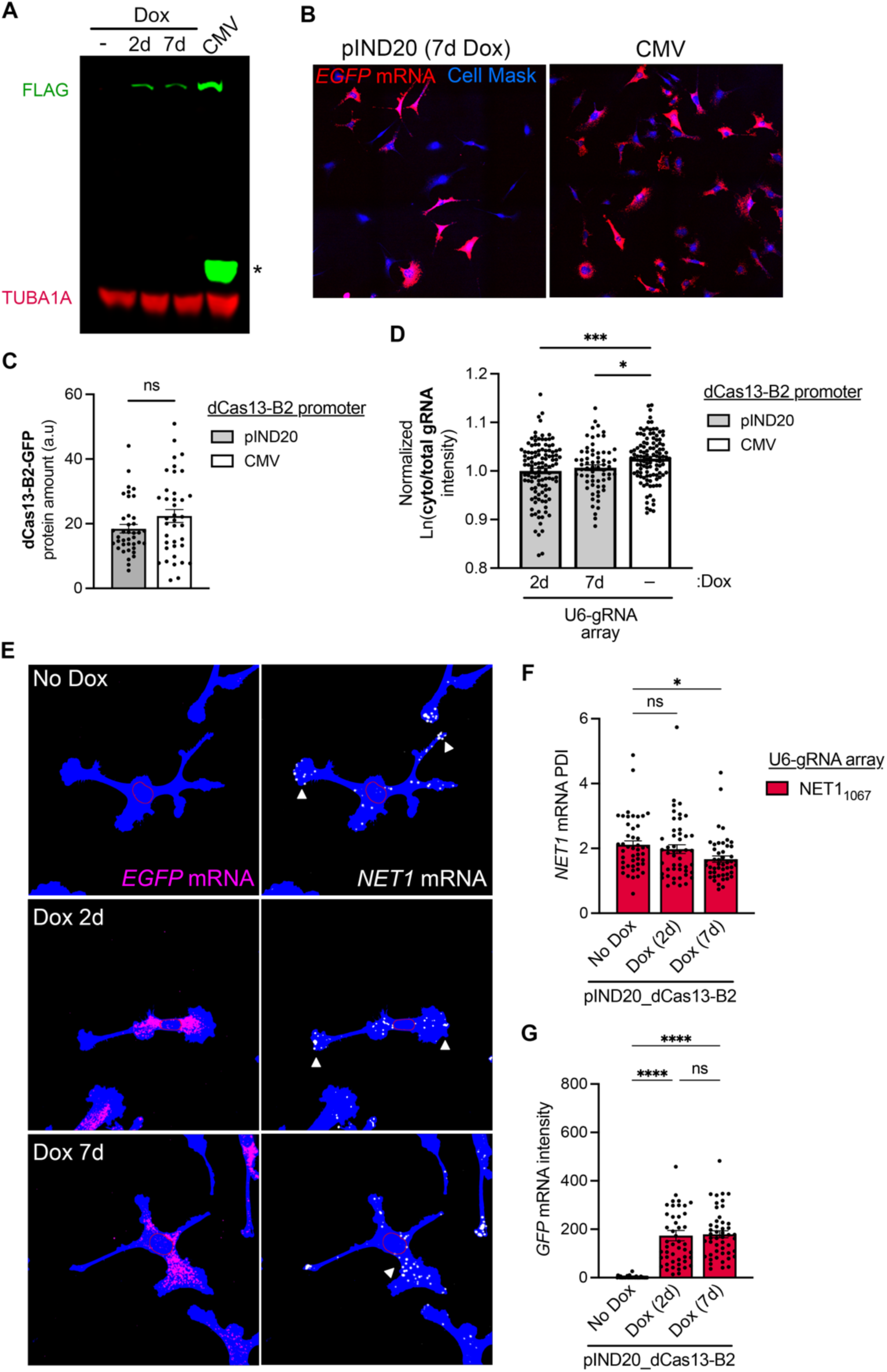
Prolonged induction of dCas13-B2 is required to alter mRNA localization. (**A**) Representative western blot of dCas13-B2 protein (detected through FLAG tag) stably expressed under either a dox-inducible or constitutive promoter. Asterisk indicates a truncated C-terminal fragment. (**B**) Representative FISH images of *GFP* mRNA in stable cell lines. CMV_dCas13-B2 cell lines more uniformly express dCas13-B2 constructs, related to Fig. 5B. (**C**) dCas13-B2 protein amount measured by GFP immunofluorescence. n= 39 cells. n.s.: non-significant by two-tailed Student’s unpaired t-test. (**D**) Cytoplasmic/total NET1_1067_ gRNA amount (from FISH images as shown in Fig. 5C) in stable cell lines with inducible (pIND20) or constitutive (CMV) dCas13-B2 expression. Days of Dox induction are indicated. n = 65-115 cells per condition; * *P* < 0.05 and *** *P* < 0.001 by one-way ANOVA with Sidak’s multiple comparisons test. (**E**) Representative FISH images of *GFP* and *NET1* mRNA upon induction of dCas13-B2 for 2 or 7 days. Arrowheads indicate areas of *NET1* mRNA accumulation. (**F**, **G**) *NET1* mRNA PDI (F) and *GFP* mRNA abundance (G) upon dCas13-B2 induction for 2 or 7 days. n = 46-48 cells per condition; not significant (ns); * *P* < 0.05 and **** *P* < 0.0001 by one-way ANOVA with Dunnett’s multiple comparisons test.

**Supplementary Figure S7.**
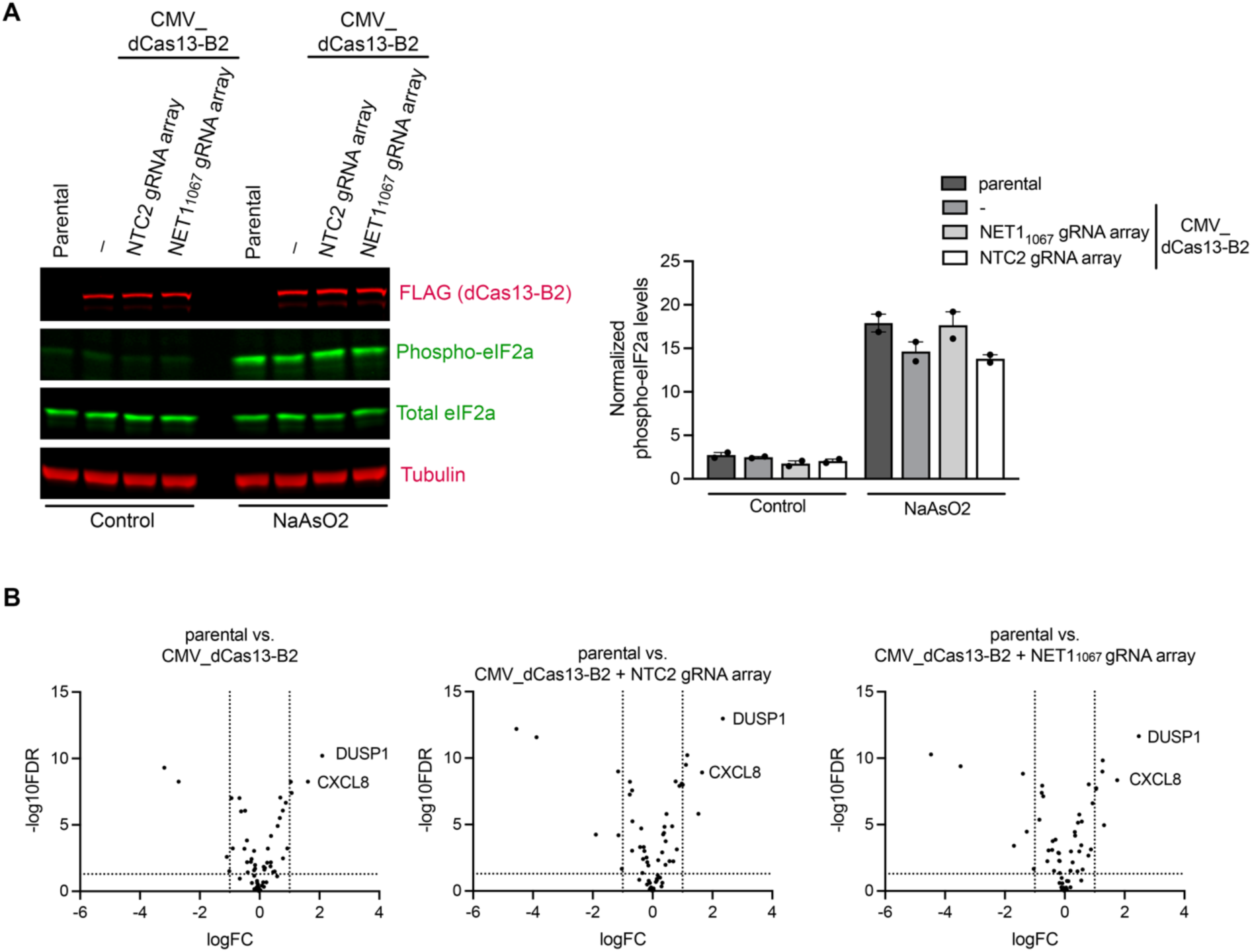
Stable expression of dCas13-B2 and gRNA arrays does not induce cellular stress response. (**A**) Western blot detection of phosphorylated eIF2α and quantification from n=2 independent experiments. (**B**) Volcano plots showing differential gene expression from the indicated cell lines. n=3 replicates per sample. Shown are 61 genes associated with cellular stress and cytotoxicity including immediate-early response, oxidative stress, ER stress, DNA damage, and apoptosis-associated genes. The genes were selected based on established stress-response pathways and Hallmark gene-set annotations and a full list and source data are shown in Supplementary Table S4. Dashed lines indicate fold change>2 and p-value<0.05. Note that the majority of genes do not exhibit altered expression indicating the absence of an overt stress response compared to the parental cell population. The consistent upregulation of DUSP1 and CXCL8 is currently of unknown significance.

**Table S1:**
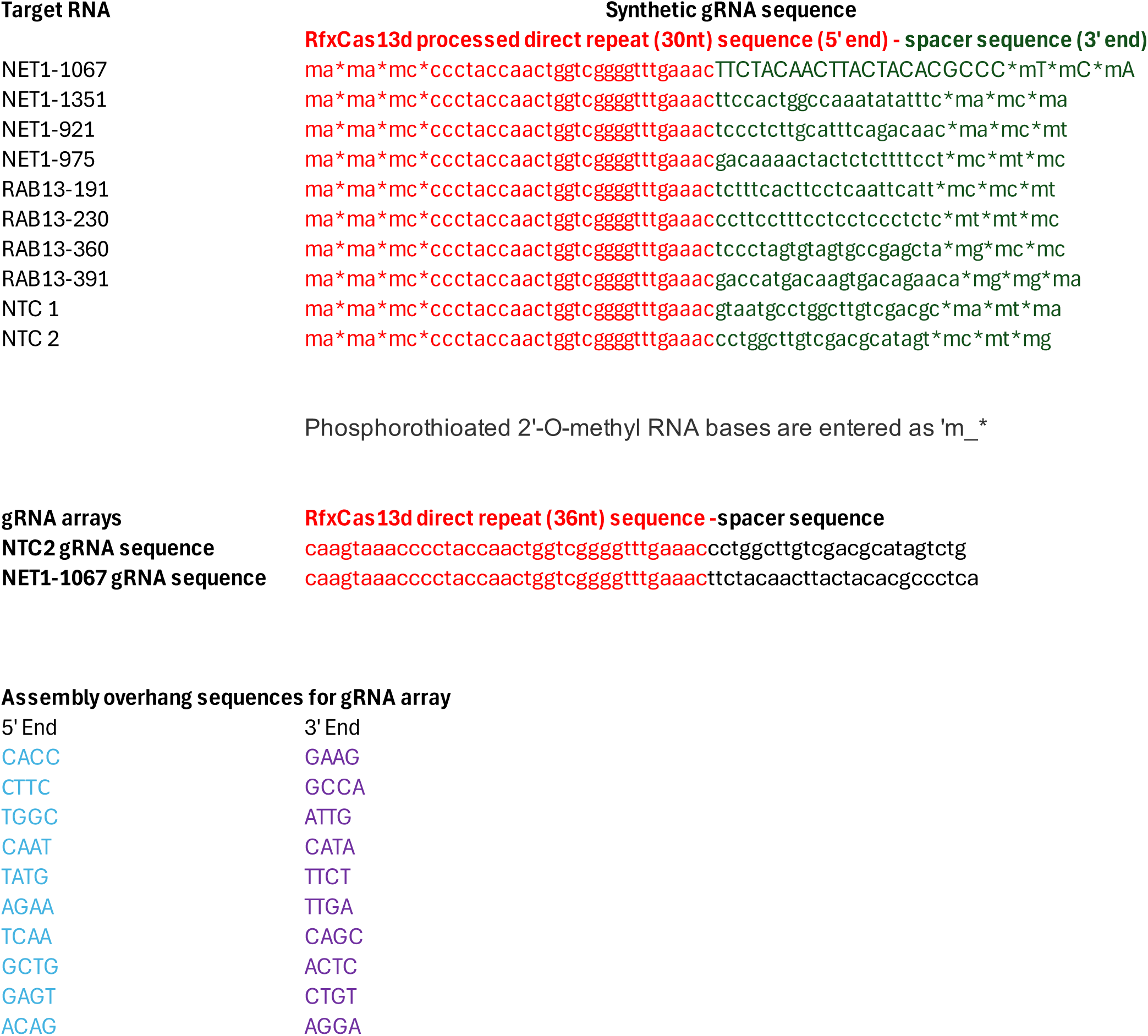
gRNA sequences.

**Table S2:**
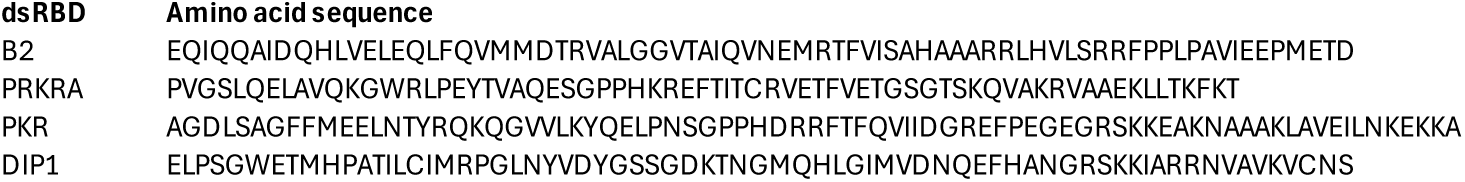
Amino acid sequences of dsRBD domains.

**Table S3:**
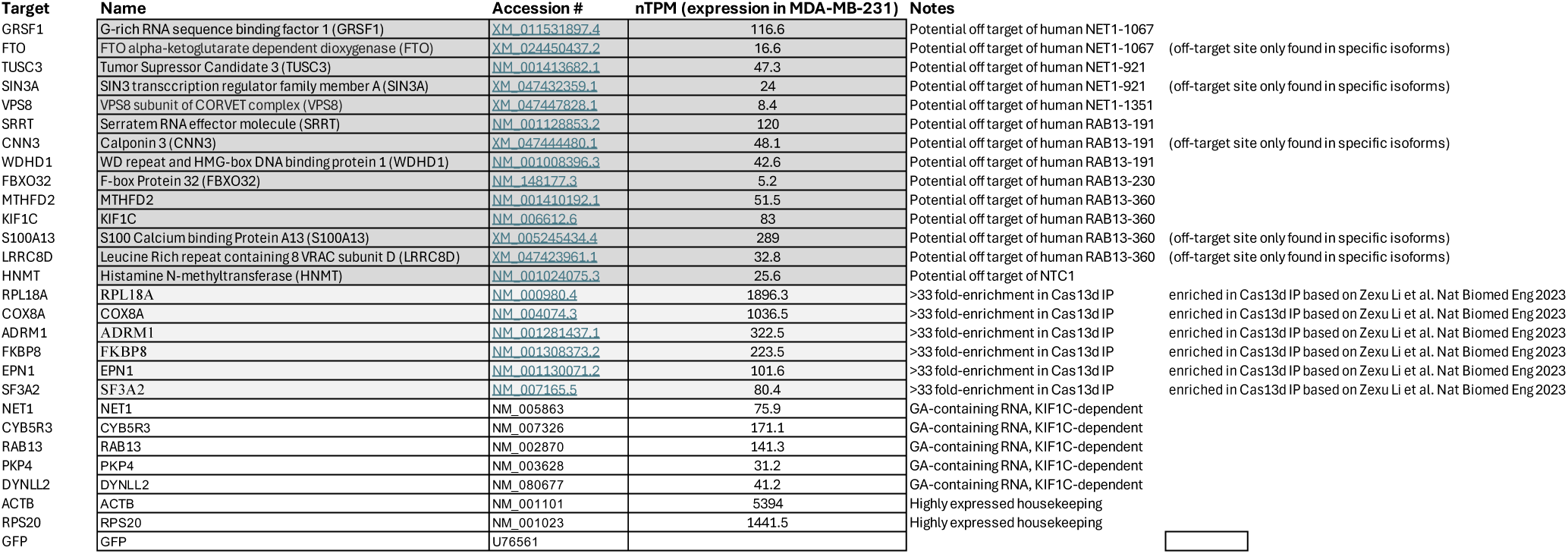
nanoString codeset targets and source data for Fig 2C-F.

**Table S3:**
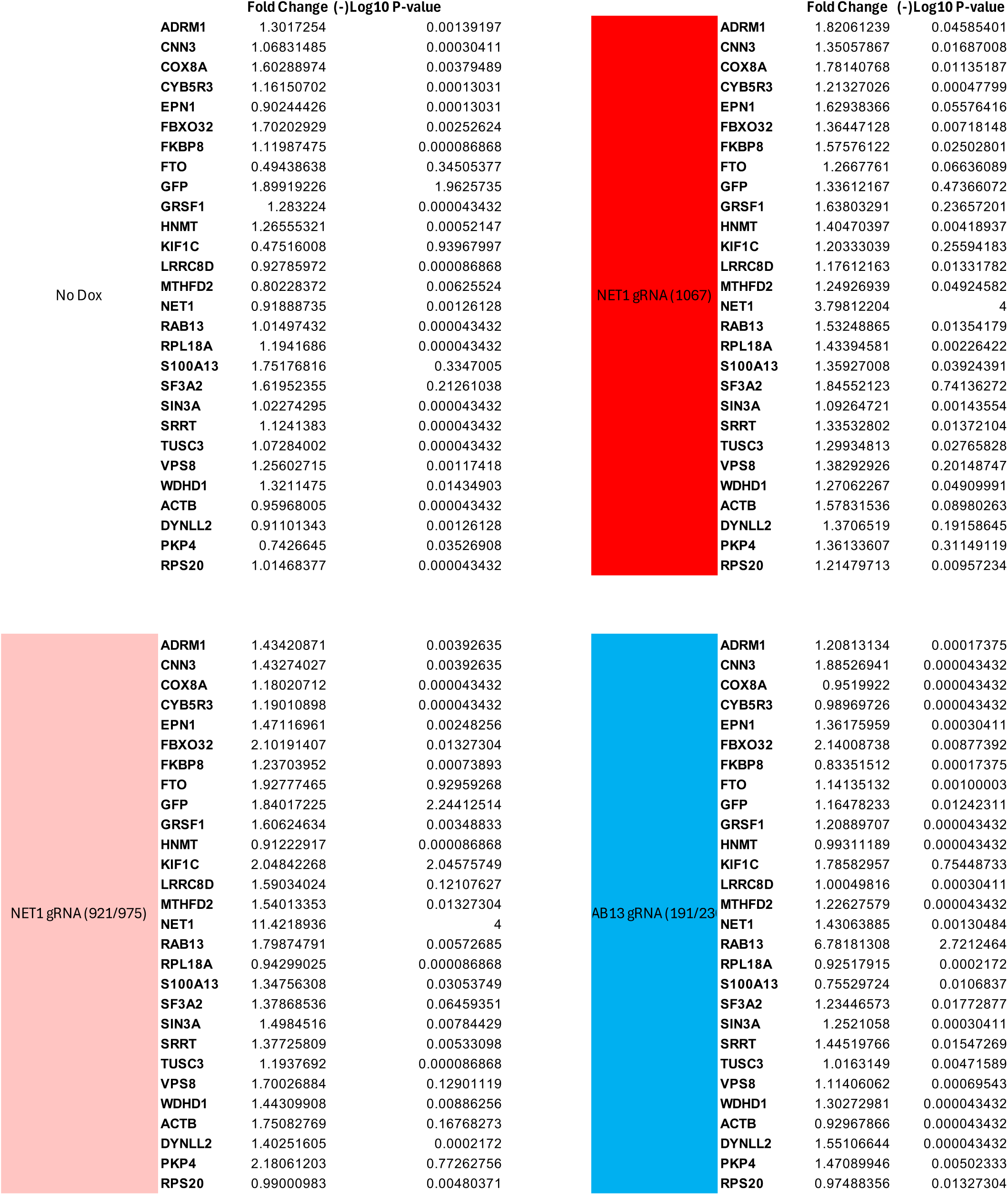
(Source data for Fig 2C-F)

**Table S4:** Source data for Supplementary Fig S7B.

| gene_name | parental vs.<br>CMV_dCas13-B2 |  | parental vs.<br>CMV_dCas13-B2 + NTC2 gRNA<br>array |  | parental vs.<br>CMV_dCas13-B2 + NET11067 gRNA<br>array |  |
| --- | --- | --- | --- | --- | --- | --- |
|  | logFC | log10FDR | logFC | log10FDR | logFC | log10FDR |
| AKR1C1 | -3.18378 | 9.30659 | -4.55264 | 12.199 | -4.46684 | 10.2797 |
| AKR1C2 | -2.71073 | 8.2614 | -3.88144 | 11.5636 | -3.48201 | 9.38988 |
| ASNS | 0.22339 | 0.681019 | -0.068849 | 0.149247 | -0.114296 | 0.278341 |
| ATF3 | 0.77832 | 2.48057 | 0.690092 | 2.23575 | 0.793903 | 2.66899 |
| ATF4 | 0.691352 | 7.05943 | 0.767333 | 8.24939 | 0.801318 | 8.03244 |
| BAG3 | -0.269129 | 2.42897 | 0.0225699 | 0.0847313 | -0.165442 | 1.36113 |
| BAX | -0.176099 | 0.173131 | 0.299777 | 0.341714 | 0.554357 | 0.797743 |
| BBC3 | -0.663715 | 0.955015 | -0.0958376 | 0.114515 | 0.192453 | 0.283127 |
| BCL2L11 | -0.419147 | 2.18936 | -0.342356 | 1.35696 | -0.541662 | 3.05378 |
| BTG2 | -1.09445 | 2.59979 | -1.8946 | 4.23768 | -1.69907 | 3.40748 |
| CASP3 | 1.04101 | 8.24863 | 1.15749 | 10.2158 | 1.26938 | 9.82747 |
| CASP7 | -0.0113553 | 0.030292 | 0.110862 | 0.681899 | 0.193362 | 1.52542 |
| CASP8 | -0.00932343 | 0.0284792 | -0.13206 | 0.763852 | -0.0254565 | 0.0994377 |
| CASP9 | -0.398651 | 1.43281 | -0.688245 | 3.03185 | -0.356179 | 1.52374 |
| CDKN1A | -0.510735 | 3.21103 | -0.67455 | 5.24621 | -0.8477 | 5.36762 |
| CXCL8 | 1.61139 | 8.25308 | 1.65208 | 8.92371 | 1.75325 | 8.34838 |
| DDIT3 | 0.874299 | 6.66493 | 0.895436 | 7.90756 | 0.932622 | 6.60596 |
| DNAJB1 | -0.185726 | 1.57952 | -0.0521091 | 0.26956 | -0.0819707 | 0.5777 |
| DNAJB9 | 0.119508 | 0.681019 | 0.167732 | 1.19495 | 0.192143 | 1.51899 |
| DUSP1 | 2.08947 | 10.2018 | 2.34593 | 12.9652 | 2.47584 | 11.657 |
| EGR1 | 0.585207 | 1.13475 | 0.324277 | 0.622937 | 0.595072 | 1.61807 |
| FAS | -0.898939 | 3.23237 | -1.1407 | 4.19778 | -1.27086 | 4.46273 |
| FOS | -0.031929 | 0.0361698 | -0.0329976 | 0.046616 | -0.0311218 | 0.0436716 |
| FTH1 | -0.18574 | 3.04498 | -0.143112 | 1.92298 | -0.192369 | 2.84547 |
| FTL | -0.0294326 | 0.274859 | -0.296677 | 3.30335 | -0.185979 | 2.28265 |
| GADD45A | 0.27317 | 3.22036 | 0.378453 | 4.36427 | 0.488743 | 5.75413 |
| GADD45B | 0.77712 | 6.08174 | 0.972925 | 8.08679 | 1.05323 | 7.74931 |
| GCLC | -0.178219 | 1.16385 | -0.319988 | 2.40883 | -0.419038 | 3.09133 |
| GCLM | -0.0612194 | 0.40968 | 0.0111691 | 0.04954 | -0.00685614 | 0.0349807 |
| GDF15 | -0.0217089 | 0.0258791 | -0.448989 | 0.848215 | -0.089167 | 0.10881 |
| HERPUD1 | 0.0249882 | 0.09942 | 0.143285 | 0.802753 | 0.12004 | 0.749235 |
| HMOX1 | 0.365966 | 1.87996 | 0.431441 | 1.96999 | 0.485083 | 2.98466 |
| HSP90AA1 | 0.382201 | 4.16638 | 0.464024 | 5.79895 | 0.440947 | 5.14308 |
| HSPA1A | -0.422649 | 3.82817 | -0.0912561 | 0.606552 | -0.0454386 | 0.259479 |
| HSPA1B | -0.152702 | 1.70877 | 0.032269 | 0.189024 | 0.0758178 | 0.75349 |
| HSPA5 | -1.01442 | 1.51581 | -1.02918 | 1.66611 | -1.03388 | 1.67128 |
| HSPH1 | 0.174269 | 1.75319 | 0.35073 | 4.25254 | 0.338718 | 4.44597 |
| IER2 | -0.260307 | 0.625336 | 0.0376019 | 0.0612495 | 0.0691591 | 0.123548 |
| IER3 | 0.25517 | 2.15001 | 0.318413 | 2.90671 | 0.26386 | 2.15026 |
| IGFBP3 | -0.942986 | 7.02362 | -1.15078 | 8.99835 | -1.39822 | 8.8351 |
| IL6 | 0.924781 | 3.23517 | 1.53247 | 5.81051 | 1.31604 | 4.95394 |
| JUN | 0.159812 | 1.62591 | 0.398562 | 4.83318 | 0.34503 | 4.16549 |
| JUNB | 0.376509 | 2.18639 | 0.654618 | 4.87479 | 0.555125 | 3.44169 |
| JUND | -0.153125 | 1.98447 | -0.380631 | 4.70729 | -0.308434 | 3.87173 |
| MDM2 | 0.603332 | 4.91572 | 0.434195 | 3.74528 | 0.566755 | 5.23318 |
| NFE2L2 | -0.0534023 | 0.502663 | -0.177912 | 2.16486 | -0.211713 | 2.879 |
| NQO1 | -0.677279 | 7.02474 | -0.686695 | 7.5652 | -0.716953 | 7.13042 |
| PMAIP1 | -0.621612 | 6.02122 | -0.76521 | 7.25181 | -0.771931 | 7.38859 |
| PPP1R15A | 1.06864 | 7.40001 | 1.11518 | 9.49125 | 1.26033 | 8.99157 |
| RRM2B | -0.29412 | 2.21659 | -0.421323 | 3.31054 | -0.400273 | 3.75064 |
| SERPINE1 | 0.137213 | 1.8482 | 0.187716 | 2.31442 | 0.209304 | 2.97968 |
| SESN1 | -0.125285 | 0.223788 | -0.184048 | 0.488152 | -0.578769 | 2.24875 |
| SESN2 | 0.450716 | 1.45757 | 0.566139 | 2.24225 | 0.425729 | 1.48473 |
| SLC7A11 | 0.67767 | 5.52245 | 1.00803 | 7.99951 | 1.0484 | 7.68701 |
| SRXN1 | -0.10048 | 0.281753 | -0.0741546 | 0.208351 | -0.0102842 | 0.0237778 |
| TNFRSF10B | 0.0778507 | 0.428297 | 0.14238 | 0.908546 | 0.0471874 | 0.250161 |
| TP53 | -0.0805478 | 0.632188 | -0.275235 | 3.00371 | -0.104565 | 0.982251 |
| TRIB3 | 0.502185 | 1.49781 | 0.80626 | 3.1308 | 0.876881 | 3.13524 |
| TXNRD1 | -0.48647 | 6.06719 | -0.75978 | 8.22312 | -0.750382 | 7.93409 |
| XBP1 | -0.0845199 | 0.777274 | -0.205124 | 2.52147 | -0.169035 | 2.26065 |
| ZFP36 | 0.140622 | 0.347063 | 0.244456 | 1.00615 | -0.177332 | 0.602743 |

